# Pseudo-dynamic analysis of heart tube formation in the mouse reveals strong regional variability and early left-right asymmetry

**DOI:** 10.1101/2021.10.07.463475

**Authors:** Isaac Esteban, Patrick Schmidt, Susana Temiño, Leif Kobbelt, Miguel Torres

## Abstract

Understanding organ morphogenesis requires a precise geometrical description of the tissues involved in the process. In highly regulative embryos, like those of mammals, morphological variability hinders the quantitative analysis of morphogenesis. In particular, the study of early heart development in mammals remains a challenging problem, due to imaging limitations and innate complexity. Around embryonic day 7.5 (E7.5), the cardiac crescent folds in an intricate and coordinated manner to produce a pumping linear heart tube at E8.25, followed by heart looping at E8.5. In this work we provide a complete morphological description of this process based on detailed imaging of a temporally dense collection of embryonic heart morphologies. We apply new approaches for morphometric staging and quantification of local morphological variations between specimens at the same stage. We identify hot spots of regionalized variability and identify left-right asymmetry in the inflow region starting at the late cardiac crescent stage, which represents the earliest signs of organ left-right asymmetry in the mammalian embryo. Finally, we generate a 3D+t digital model that provides a framework suitable for co-representation of data from different sources and for the computer modelling of the process.

**SUMMARY STATEMENT:** We provide the first complete atlas for morphometric analysis and visualization of heart tube morphogenesis, reporting morphological variability and early emergence of left-right asymmetry patterns.

## INTRODUCTION

Quantitative morphometric analysis is required for a deep understanding of the mechanisms that drive embryonic morphogenesis. Defining accurate geometries for highly dynamic morphogenetic processes with inherent variability represents an important challenge that remains unresolved. The mammalian embryo is highly regulative, which implies it can consistently develop stereotyped organs despite initial morphological heterogeneity. Heart tube (HT) formation involves complex morphogenetic movements and differentiation patterns, through which form and function are acquired simultaneously (Kelly et al., 2014). Understanding heart tube morphogenesis would greatly benefit from having access to a spatiotemporal representation of the morphology of the tissues involved in the process. Previous attempts to understand the 3D+t complexity of primitive heart tube formation in amniotes have been based on the segmentation and 3D reconstruction of specimens from tissue sections (de Bakker et al., 2016; de Boer et al., 2012; Faber et al., 2021; Soufan et al., 2006). This approach has allowed the reconstruction a few discrete stages capturing this process, but is limited by a demanding methodology and the distortions of section alignment. Additional efforts have included high-resolution episcopic microscopy and live analysis of the developing chicken or mouse heart; however, this was applied to stages beyond primitive heart tube formation and focused on specific processes of cardiogenesis (Kawahira et al., 2020; Lopez et al., 2020; Mohun and Anderson, 2020; Yue et al., 2020) or was limited by incomplete imaging of the area of interest and low image resolution (Ivanovitch et al., 2017; Le Garrec et al., 2017).

Cardiac development starts with the deployment of cardiac mesoderm early in gastrulation (Meilhac and Buckingham, 2018). Soon after colonizing the antero-lateral region of the embryo, the cardiac mesoderm at the anterior lateral plate undergoes a mesenchymal to epithelial transition to produce two layers that enclose the celomic cavity. This results in two independent epithelial sheets: the splanchnic mesoderm, which shares basement membrane with the endoderm, and the somatic mesoderm which shares basement membrane with the ectoderm. The two mesodermal layers meet medially with the head paraxial mesoderm and laterally at the embryonic-extraembryonic border, forming a closed and continuous three-dimensional surface encasing the anterior celomic cavity (future pericardial cavity).

During the E7.5-E8.5 developmental window, the splanchnic and somatic mesoderm layers progressively move apart from each other, forming the pericardial cavity, which spans from the antero-medial region to the lateral sides. This occurs in parallel to embryonic folding and foregut pocket formation, which brings the foregut endoderm to the inside of the embryo. This causes the pericardial cavity to rotate inwards around the left-right axis, driven by the rostral folding of the embryo, concomitant with the invagination of the endoderm towards the anterior-dorsal region. Following foregut pocket formation, the cavity is limited at its cranial side by the ectodermal plate and the head mesoderm. Caudally, the cavity limits with the foregut diverticulum in its medial aspect, while the posterior-lateral limits of the cavity extend around the foregut invagination, ending at the somatic and splanchnic mesoderm junction.

The splanchnic mesoderm of the forming pericardial cavity contains the cardiac mesoderm that gives rise to the HT. Within the cardiac mesoderm, there are two distinct regions; the first heart field (FHF), which differentiates rapidly to the cardiomyocyte fate, and the second heart field (SHF), which remains highly proliferative and undifferentiated until its precursors are progressively incorporated to the heart at later stages. While the FHF is underlined by endothelial cells (precursors of the endocardium), the SHF remains directly attached to the basement membrane shared with the endoderm. The FHF extends antero-laterally, closer to the embryonic-extra-embryonic boundary, while the SHF occupies the dorso-medial regions adjacent to the FHF. The initially relatively flat cardiac crescent (CC) starts very active morphogenesis around E7.5, growing and acquiring three-dimensional complexity by folding on itself, detaching from the endoderm and coalescing medially to form the primitive tube, with the venous to arterial pole axis in a caudal to cranial orientation.

In this work, we aimed to obtain a complete description of tissue anatomy during primitive heart tube morphogenesis in the mouse embryo. To this end, we acquired high-resolution confocal images at high temporal density from whole specimens collected between E7.5 and E8.8. We used this image collection to elaborate a continuous 3D+t description of heart tube morphogenesis through a new morphometric staging system and a 3D interpolation strategy. Given that heart tube morphogenesis cannot be uncoupled from the morphological evolution of the surrounding tissues, our model incorporates all pericardial cavity mesoderm, endoderm and the endothelium. Besides obtaining a detailed 3D+t atlas of heart tube formation and its associated tissues, we applied statistical methods to study regional variability and left-right patterning during heart tube formation. The model and knowledge generated provide a strong basis for understanding heart morphogenesis, its variability and its congenital alterations.

## RESULTS

### Acquisition of high-resolution images of the developing heart at high temporal resolution

To generate a high-fidelity 3D+t atlas of early mouse heart development, we imaged 52 fixed and cleared (Susaki et al., 2014) embryos in cardiac developmental stages ranging from the early CC until heart looping. Embryos were collected at nominal stages from E8 to E8.25, however, given the variability of stages found within littermates, the collection represents nominal ages from approximately E7.75 to E8.5 (18 hours). The estimated approximate temporal density is therefore 1 specimen every 20 minutes. The imaged embryos carried *Mesp1*^*Cre*^ and *R26R*^*mTmG*^ and *R26R*^*Tomato*^ alleles, so that all mesoderm in the head and cardiogenic area was labelled with membrane-GFP and cytoplasmic Tomato, while the rest of the tissues were labelled only with membrane-Tomato. We imaged volumes of ~1mm in the X and Y dimension and down to a depth of 677µm, so that the whole spatial domain of the cardiogenic region and associated tissues was acquired. In order to obtain high-resolution images suitable for precise 3D segmentation, we used an XY pixel resolution varying from 0.38µm to 0.49µm and a z-step varying from 0.49µm to 2.0µm, depending on the stage. This strategy provided highly detailed orthogonal optical sections (Fig. 1C, see materials and methods).

**Figure 1.**
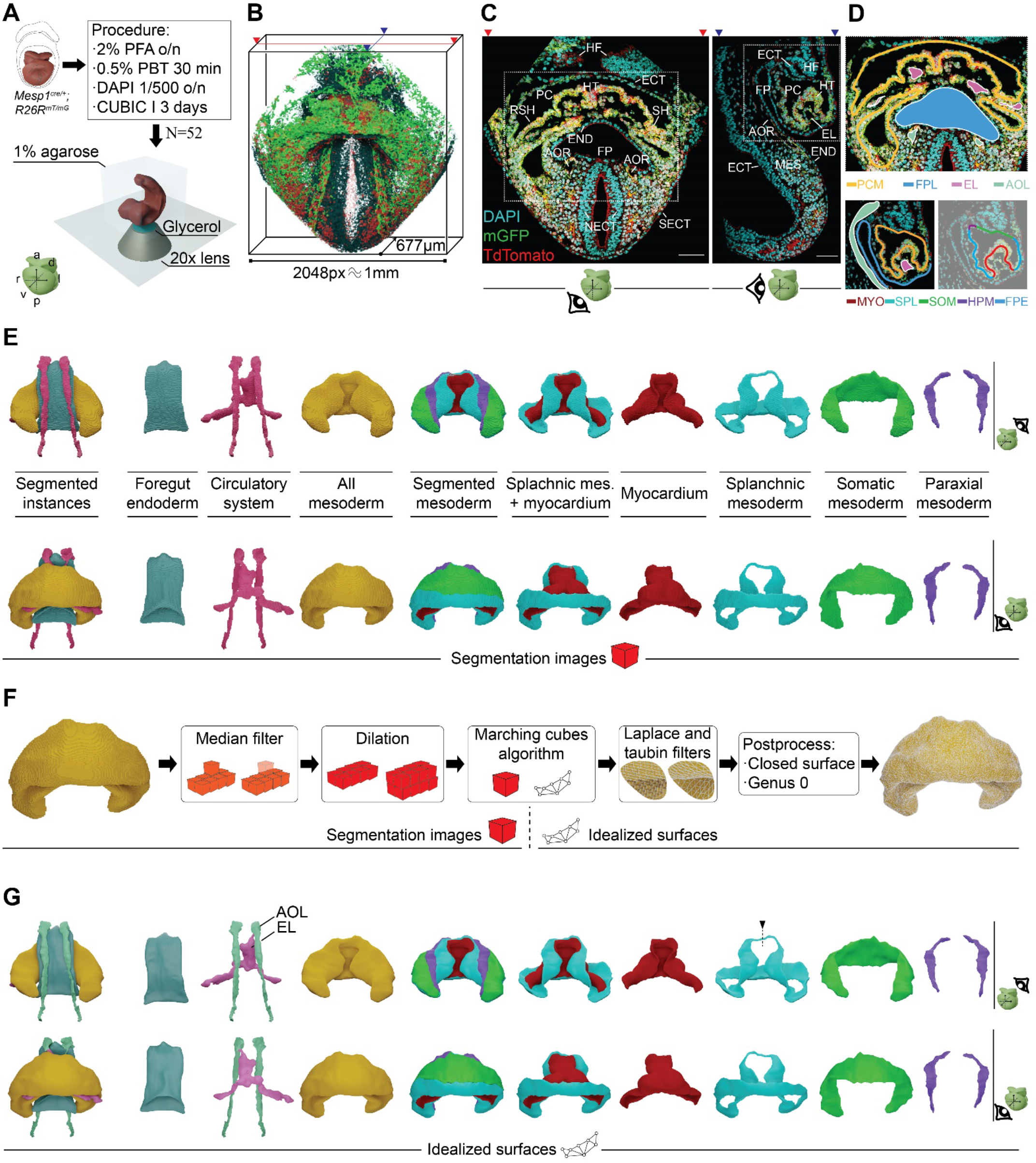
Segmentation of high-resolution confocal images reveals precise and detailed morphological representations for understanding heart development. (A) Schematic representation of the workflow for image acquisition. Below is represented the axis of the system of reference. (B) Confocal raw image reconstruction showing the extension of the imaged spatial domain. Mesodermal tissues show the cytoplasm in red and cell membranes in green; the rest of tissues show membranes in red; cell nuclei appear in cyan. The acquisition window resolution and maximum depth reached are shown. Red arrows show the direction of the imaging plane and blue arrows the direction of the lateral orthogonal projection. (C) (Left) Frontal optical section (corresponding to red arrows in B) captured using confocal microscopy. Dotted-line box illustrates the extent of the cardiac region. (Right) Sagittal optical section (corresponding to blue arrows in B). High resolution at both the imaging plane and the orthogonal views allows for a precise segmentation. Scale bar 100µm. (D) (Top) Frontal optical section at the level of the cardiac inflow region; the orange line represents the mesoderm of the anterior celomic cavity, including the differentiating myocardium; the foregut pocket lumen is represented in blue; the endocardial lumen is represented in pink, and the aortic lumen is represented in green. (Bottom left) shows a lateral sagittal optical section at the level of one of the aortas. (Bottom right) the same optical section showing the subtypes of mesoderm of the pericardial cavity. Myocardium in red, splanchnic mesoderm in blue, somatic mesoderm in green and head paraxial mesoderm in purple. (E) 3D representation of the segmented images. Top and bottom rows show dorsal and ventral views, respectively. (F) Pipeline for transforming the segmentation images into smooth, closed meshes of genus 0. (G) Resulting meshes. The circulatory system is split in two parts: the lumen formed by the endocardial cells and the aortas. Top and bottom rows show dorsal and ventral views, respectively. The black dotted-line and arrow represent the cut trajectory to achieve genus 0 topology in the splanchnic shapes (see material and methods). **HF**, head folds; **HT**, heart tube; **ECT**, ectoderm; **PC**, pericardial cavity; **RSH**, right sinus horn; **LSH**, left sinus horn; **END**, endoderm; **FP**, foregut pocket; **AOR**, aorta; **SECT**, surface ectoderm; **NECT**, neuro-ectoderm; **EL**, endocardial lumen; **MES**, mesoderm; **PCM**, pericardial cavity mesoderm; **FPL**, foregut pocket lumen; **FPE**, foregut pocket endoderm; **AOL**, aortic lumen; **MYO**, myocardium; **SPL**, splanchnic mesoderm; **SOM**, somatic mesoderm; H**PM**, head paraxial mesoderm. Specimen shown in 1A-1D corresponds to E32; specimen shown in 1E-1G corresponds to E40.

We then semi-automatically segmented (Yushkevich et al., 2006) the mesoderm of the anterior celomic cavity, discriminating both layers of the lateral plate mesoderm; splanchnic and somatic, up to their fusion with the head paraxial mesoderm (Fig. 1D). Within the splanchnic domain, we used morphological and topological criteria to discriminate the differentiated myocardium from the rest of the splanchnic mesoderm. To do so, the cells that were found detached from the endoderm and encasing or overlying endocardial cells were classified as differentiating cardiomyocytes (Fig. 1D). In addition, we fully automatically (He et al., 2017) segmented the foregut endoderm and the forming aortae. To reconstruct the incipient circulatory system, we joined the automatically segmented aortae with the semi-automatically segmented endocardial cavity (Fig. 1D). The segmentation was done in each 2D optical section from the confocal images, combining the information from the three orthogonal views to increase segmentation accuracy, and then rendered in 3D (Fig. 1E). The segmentation images were then subjected to filtering (Cignoni et al., 2008) in order to remove noisy voxels (Fig. 1F). Finally, the marching cubes algorithm (Lorensen and Cline, 1987) was applied to obtain a discrete numerical description of the surface of each tissue in the form of a mesh (Fig. 1F), followed by smoothing filters. Post-processing steps were then taken to ensure closed, oriented, genus-0 surface meshes of the tissue (Fig. 1F), for which we used Version 2.37.13 of Trimesh (2019). In this way, we generated 3D models representing the segmented cardiac tissues and the surrounding tissues for the 52 specimens of the collection (an example is shown in Fig. 1G). These 3D models are available for download and can, for example, be visualized using the open-source application ParaView.

Preliminary study of 5 selected specimens in obvious temporal sequence (Fig. 2A, B), showed how the splanchnic mesoderm folds to form the CC. The early CC extends bilaterally and subdivides the undifferentiated splanchnic mesoderm into medial and distal domains. The medial domain contains the SHF precursors, while the distal domain (recently named juxta-cardiac field) contains precursors of the epicardium and the *septum transversum*, and bears myocardial differentiation capacity (Tyser et al., 2021). As the cardiac crescent undergoes morphogenesis to generate the cardiac tube, both the juxta-cardiac field and its border with the forming HT remain morphologically rather stable (Fig. 2A, B), whereas the medial splanchnic mesoderm (containing the SHF) and its border with the forming HT undergo a drastic deformation that is essential for the generation of the outflow tract and the dorsal closure, which transforms the CC into the primitive HT (Fig. 2A, B). This transformation is concomitant with the development of the endocardial cavity, the aortae, and their connection through the first branchial arch arteries (Fig. 2C, D).

**Figure 2.**
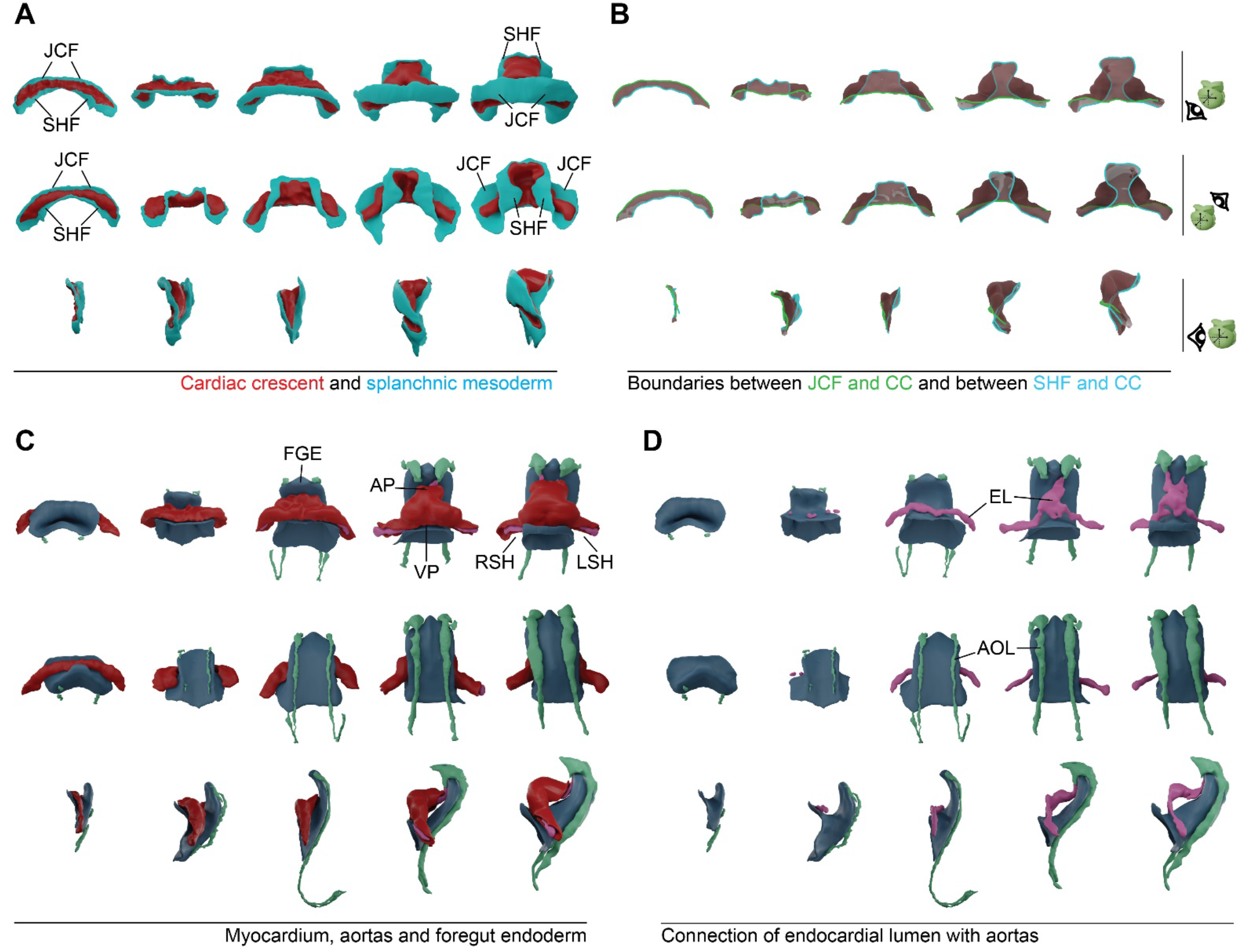
Observation of qualitative morphometric aspects of heart tube morphogenesis. (A) Ventral, dorsal and lateral views of 5 specimens, subjectively ordered according to developmental time. Splanchnic mesoderm is represented in blue and differentiated myocardium in red. The dorsal view shows the progressive dorsal closure of the heart tube and the medial expansion of the SHF. (B) Limits of the myocardial differentiated domain, depicted on the edges of the myocardium mid-surface. The green line represents the boundary between the juxta-cardiac field and the myocardium. The blue line represents the boundary between the myocardium and the SHF. (C) Morphology of the myocardium together with the foregut endoderm and the circulatory system lumen. The different length of the aortae seen in the three older specimens, is due to variable imaging depth. (D) Gradual development of the circulatory system, in parallel to the foregut endoderm invagination. Ventral view shows the progressive formation of the endoderm lumen. Lateral view shows the progressive formation of the aortic lumen in along the dorsal surface of the foregut endoderm. **JCF** juxta-cardiac field; **SHF**, second heart field; **FGE**, foregut endoderm; **AP**, arterial pole; **VP**, venous pole; **EL**, endocardial lumen; **AOL**, aortic lumen; **LSH**, left sinus horn; **RSH**, right sinus horn. Specimens shown in the figure are E1, E7, E11, E27 and E39.

### Development of a morphometric staging system for heart tube formation

We next aimed for an accurate temporal ordering of all the specimens rendered. The examination of the collection of datasets reflected high morphological heterogeneity of the developing heart within groups of embryos at an apparently similar developmental stage, as judged by somite number, head folds and foregut pocket shapes. The morphological heterogeneity affects both general conformation features, like proportions, degree of dorsal closure or degree of looping, and local morphological aspects, such as non-stereotyped presence of bulges of different sizes and distribution. This suggested that it is very complicated to define a staging system based on simple morphological criteria. In order to tackle the problem of staging, we then implemented different morphometric strategies to analyse the 3D reconstructions. We explored the measurement of different landmark curves and distances defined on the surface of the tissues, seeking for simple parameters that could report the temporal evolution of the shapes. For this purpose, we first obtained mid-surfaces (Yoshizawa et al., 2003) for each of the relevant tissues, consisting of a zero-thickness surface resulting from flattening the original surfaces (Fig. 3A).

**Figure 3.**
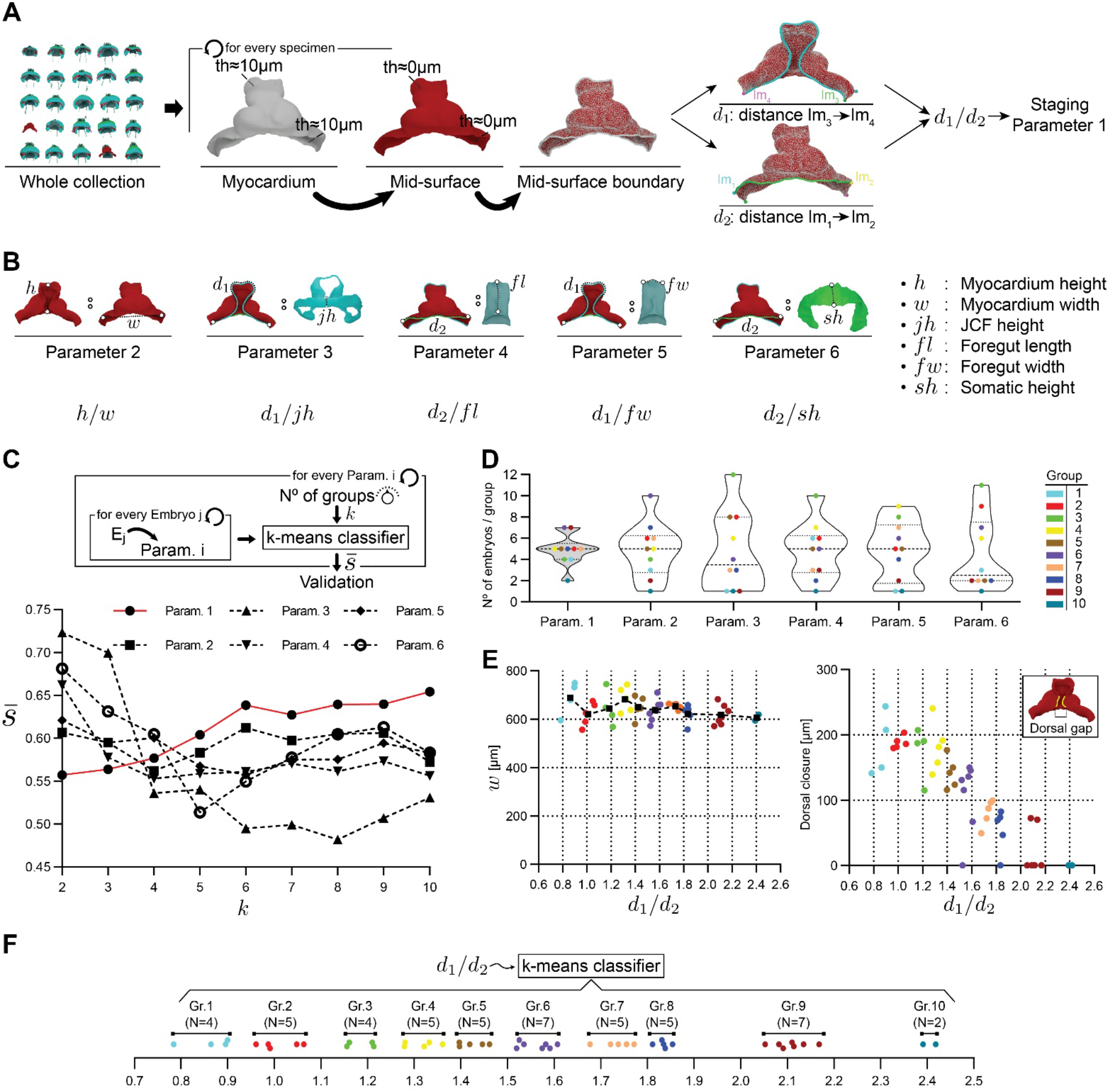
Establishment of a morphometric staging system. (A) Schematic representation for the calculation of staging parameter 1. (B) Schematic representation of five additional staging parameters. (C) (Top) Workflow for evaluating staging parameters, by means of the average silhouette coefficient (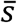). (Bottom) Graph showing the evolution of 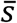 as a function of the number of groups, for every staging parameter. (D) Violin plots showing the distribution of specimens in each cluster (*N*=10 clusters) for the 5 selected staging parameters. (E) (Left) Scatter plot showing myocardium width, as a function of the staging parameter 1. (Right) Plot illustrates the dorsal gap distance, excluding early stages in which it could not be measured, because the process of dorsal closure had not initiated. (F) Plot showing the final classification of the collection in 10 groups according to parameter 1. Specimen shown in A and B is E40.

We next studied the proportions between the lengths of pairs of reference curves/distances, focusing on proportions expected to vary continuously during the studied period (See example in Fig. 3A). Given that the cardiogenic area evolves from a crescent extending from left to right to a tube extending from cranial to caudal, a general trend during the formation of the heart tube is the increase of the height (cranio-caudal size) to width (left-right size) ratio of the tissues involved. We therefore calculated the proportions between different parameters related to the height-to-width ratio variation (Fig. 3A and B, parameters 1 to 6). We then applied the *k-means* clustering algorithm for the determination of groups based on differences in the defined parameters and calculated the *average silhouette coefficient* 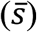 (Fig. S1), which measures the quality of classifications (Rousseeuw, 1987). As we increased the number of groups (*k*) in the k-means classification, the best 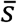were obtained for the *d1*/*d2* proportion, *d1* and d2 being the length of the border between the myocardium and JCF, and between the myocardium and SHF respectively (Fig. 3C, curves 1 and 2 in Fig. S11.). The maximum 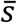was obtained for a *k-means* classification into 10 groups. For *k*>10, *s* shows oscillatory behaviour, followed by a rapid decay (not shown). We therefore concluded that the d1 to d2 ratio was the best single parameter for classifying the collection in different stages, and established these 10 groups as staging reference (Fig. 3E). Study of the correlation between d1/d2 and the rest of the parameters provided a view of how good the different classification methods are with respect to d1/d2 (Fig. S1). The best correlate to d1/d2 was the height to width ratio, which showed a high linear correlation (Fig. S2A). In contrast, other parameters either showed a poor correlation or did not show a linear correlation, which resulted in poor classification of the older specimens (Fig. S2). Two specimens (E51 and E52) were left out of the classification because they represented outliers too early or too advanced with respect to the rest of specimens (Figs. S3-S10) and one specimen (E3) in which differentiation of the cardiac crescent had started, but was not sufficient to calculate d1/d2, was manually assigned to Group 1 by similarity. The d1/d2 proportion also provided the most homogeneous distribution of specimens among the 10 stage groups, with seven of the groups containing 4 or 5 specimens, only one containing 2 and two containing 7 specimens, while all other classification methods showed strong heterogeneity in the distribution of specimens per group (Fig. 3D).

The geometries of all the tissues segmented for every specimen classified by stage are shown in Figs. S3-S10. Groups 1 to 4 represent different stages of cardiac crescent development, while Groups 5 to 8 can be assigned to linear heart tube stages, and Groups 9 and 10 to heart looping stages.

Interestingly, the absolute width of the forming cardiac tube, which coincides with the width of the pericardial cavity, not only does not increase during development, but shows a mild trend to reduction as development progresses (Fig. 3E). This aspect indicates that, despite overall embryo growth, the lateral expansion of the pericardial cavity seems restricted. In contrast, the dorso-ventral and cranio-caudal dimensions of the pericardial cavity and all associated tissues, including the forming heart tube, strongly increase during this period. This aspect correlates well with the extensive deformation that the curve d1 undergoes, associated to myocardial and splanchnic mesoderm remodelling and foregut pocket extension, in contrast with d2, which remains mostly stable. We also studied the occurrence of developmental events, like dorsal closure and looping, which are known to progress continuously in live-imaged single specimens (Ivanovitch et al., 2017; Le Garrec et al., 2017). Looping is appreciable in all specimens of group 10 and most specimens of group 9, without signs of looping in specimens of other groups, which suggests that looping starts approximately at the same maturation stage in all specimens (Figs. S5, S6). A possible exception to this is specimen E33, which appears to be starting looping, however the looping direction appears inverted in this specimen and therefore it might be a case of aberrant development. We also measured the dorsal mesocardial gap (Fig. S11A), as this gap is initially wide open in the cardiac crescent and closes until the gap disappears and the dorsal mesocardium is formed upon linear heart tube formation. While dorsal closure appeared predominantly associated with looping, and therefore was seen in all specimens in stage 10+ and most specimens in stage 9, we also found some hearts already closed at stage 6-8, while some specimens of stage 9 that were clearly advanced in looping were quite delayed in dorsal closure (Figs. S5, S6). In agreement with these observations, dorsal gap correlation with d1/d2 was quite noisy (Fig. 3E), indicating that the degree of dorsal closure (see materials and methods) is not a reliable parameter for staging.

### Definition of standard geometries for each stage of heart tube formation and mapping of local shape variability

The distribution of *d1*/*d2* values among the 10 staging groups established can be seen in Fig. 3F. Given the variability found between specimens in the same staging groups (Figs. S3-S10), we next aimed to quantify and spatially map the shape variability of the differentiated myocardium within each of the 10 stages established and to define a consensus geometry for each stage (Fig. 4A). Computing a consensus geometry within a group requires surface correspondence among them. To this end, we established a common triangulation, which allowed to compute the mean shape. We chose the triangulation of the geometry whose *d1*/*d2* was closest to the average *d1*/*d2* of the group (Fig. 4A’, left, central specimen: Y1). We then used the method described in (Schmidt et al., 2020) (Fig. 4A’, middle) to compute surface maps from the reference mesh to all the other meshes in the group. Each surface map transfers the vertices of the reference mesh to the respective target surface, establishing a vertex-to-vertex correspondence between all shapes of a group.

**Figure 4.**
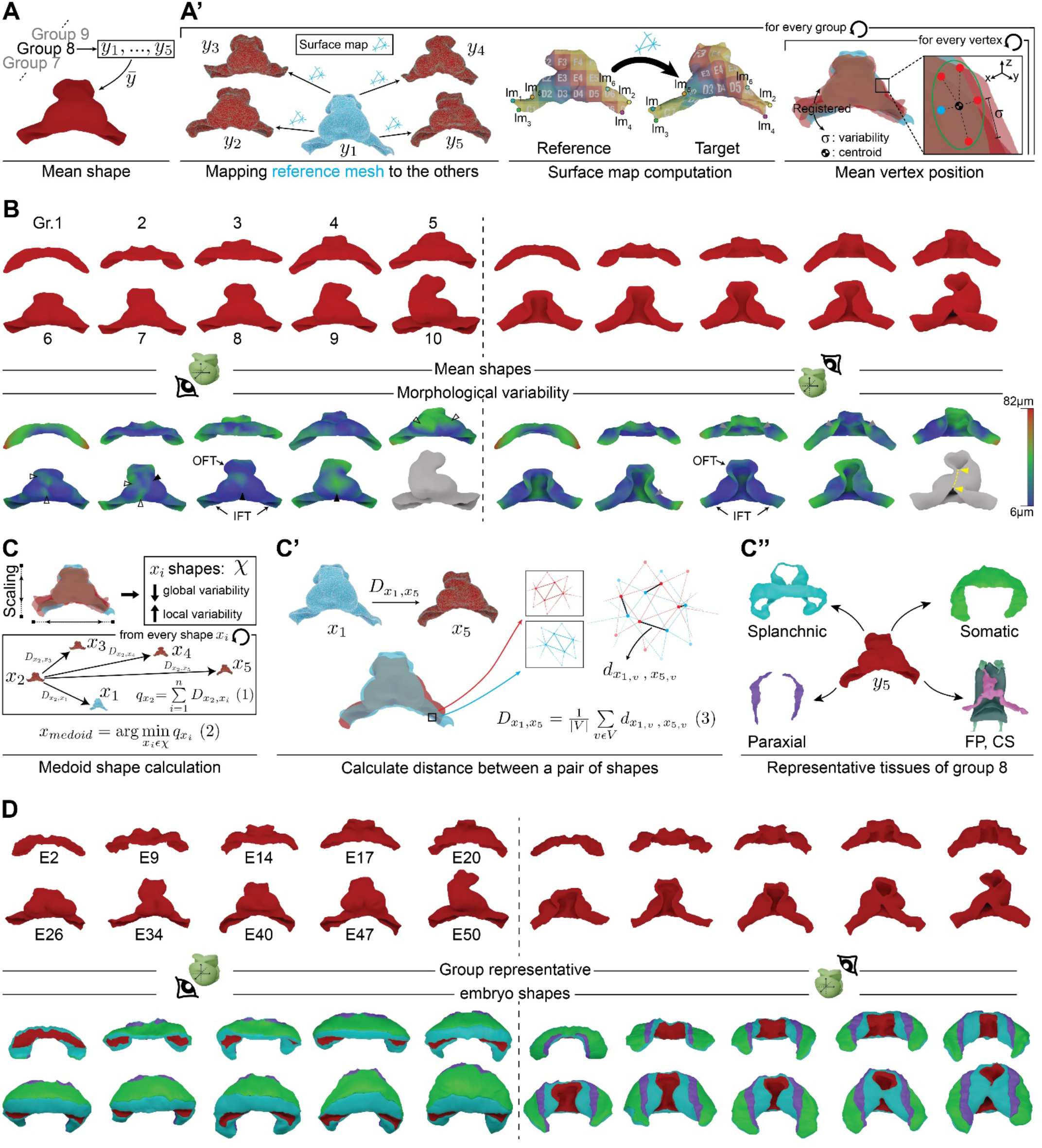
Mean shape for each group (MSG) and morphological variability. (A) Illustration of the concept of mean shape for staging group 8. (A’) Workflow for obtaining a mean shape and local variability estimation for each staging group: (Left) Process of surface map computation. The blue shape represents the mesh of the reference specimen of the group. The arrows represent the surface mapping from the reference specimen mesh to all others in the staging group. (Middle) This panel represents the working principle of the surface map computation. The landmarks (*lm*) identify corresponding locations in the reference and target mesh. (Right) Shapes of the group after rigid registration to the reference mesh. For every vertex in the reference mesh (blue dot), there exists a corresponding point on the other shapes (red dots). (B) The panel shows the calculated myocardium mean shapes, for every staging group. The top two rows show the mean shapes from ventral (left) and dorsal (right) views. The two rows below show the same mean shapes, color-coded according to the morphological variability calculated for each vertex. The shape in grey indicates that the low number of specimens at this stage precludes variability calculation. The yellow dotted-line and arrows in the dorsal view of this shape represent the cut trajectory to achieve genus 0 topology in dorsally closed hearts (see material and methods). Arrowheads indicate the regions of higher variability; solid ones indicate medial ventricular regions and open ones indicate lateral ventricular regions. (C) Workflow for calculating the medoid shape of a stage group. Rigid registration with scaling is first performed and then the term *q* is calculated for each specimen using the Eqn 1 (boxed scheme). The medoid is then calculated using Eqn 2 (C’) Graphical illustration of the calculation of *D*, as part of the Eqn 1. The blue and red meshes show an example of a zoom of the surface mesh of two shapes, in their registered position. Black lines between vertices measure the distance *d* between corresponding vertices of both shapes. Eqn 3 is used to calculate *D*. (C’’) Myocardium shape selected as the representative of group 8, together with all the associated tissues. (D) This panel shows the representative specimens of each group. Top left and right show ventral and dorsal view of the shape of the myocardium, respectively. Bottom left and right show ventral and dorsal views, respectively, of all the pericardial tissues of the representative embryos. **OFT** outflow tract; **IFT**, inflow tracts; FGE, foregut endoderm; CS, circulatory system. **y**_**i**_: specific shape of the group; **x**_**i**_: specific shape of the group after scaling; **X**: set of all x_i_ shapes; ***q***: sum of morphological distance from one shape to all others; ***d***: distance between two vertices; ***D***: average of d, calculated along the surface mesh domain; ***v***: index pointing to a specific vertex; ***V***: vertex set; ***n***: number of shapes in the group.

Each surface map computation requires a sparse set of corresponding points (landmarks) as input (Fig. 4A’, middle). We developed a methodology for defining such points (Figs. 4A’ middle, S11C). This was a challenging task, because there is no fixed reference in the embryo and only internal references could be set. We established a set of landmark points and curves that identify equivalent positions across stages, thus providing a reference for initialization of the surface map computations (Fig. S11B-E). Finally, using the dense correspondence provided by the maps, we rigidly aligned all shapes of the group to the reference shape (Fig. 4A’).

Next, we averaged the positions of corresponding vertices within the group to construct a Mean Shape for each Group (MSG) (Fig. 4A’, right). The collection of 10 MSGs therefore represents the average evolution of myocardial shape during the formation of the primitive heart tube and initiation of looping (Fig. 4B).

We then computed the per-vertex variability among the specimens of each staging group. We use this approach as a statistical framework to quantify the natural variability of heart tube morphology and to quantitatively map it in 3D. In this way we were able to identify hotspots of variability in the 10 different stages (Fig. 4B). In most groups we found that high variability appeared in the outflows, inflows and the dorsal lips of the myocardium. During primitive ventricular chamber formation, we found that variability concentrated first at the bilateral bulges that start to form the chamber and later at the midline, possibly in relationship to the variability in the degree of merging of the two initially bilateral bulges into one. Finally, in stage 9, high variability was observed in all structures involved in the looping process (Fig. 4B). Variability was not measured in stage 10, given that this group is composed of only two specimens. This collection of MSGs provides information on the standard shapes of different stages of primitive heart tube formation and the local variability of the process.

We also wanted to identify the myocardial individual shape that best represents its stage group, as this would give us access to representative geometries for the other tissues as well (Fig. 4C). For this, we calculated for each stage group the medoid shape of the group (Fig. 4C’, Equation 2) by identifying the geometry whose accumulated differences with respect to all others in the group were the smallest (Fig. 4C’’, Equation 1, Equation 3). Prior to this, all the shapes in a group are uniformly scaled (Fig. 3A’), to diminish the effect of differences in global size. In this way, we identified a concrete specimen as Stage Group Representative (SGR) for each stage (Fig. 4D).

### Morphometric comparison of left-right geometries identifies early asymmetry of the inflows

Heart tube looping is considered the first morphogenetic expression of left-right asymmetry in a developing organ. Here, we identified obvious signs of heart looping in stages 9 and 10, however we wanted to explore systematically whether less obvious asymmetries might be present before the looping stage. Studying both the whole specimen collection and the collection of MSGs, we detected a possible recurrent difference in the angle of insertion of the sinus horns (inflows). To study this aspect in more detail, we defined the direction of insertion of the inflows as a result of two polar coordinates (see materials and methods); the *θ* angle, consisting on the angle between the orthogonal projection of the inflow insertion direction on the *xy* plane and the *x* axis, and the *φ* angle, directly calculated with respect to the z axis (Fig. 5A). When studying the evolution of these angles in correlation to the staging parameter *d1*/*d2*, we found that the *θ* angle diverged between the left and right sides, while the *φ* angle remained symmetric (Fig. 5B). The *θ* angle significantly deviated from symmetry from *d1*/*d2* values of 1.6, corresponding to a staging between groups 5 and 6 (Fig. 5C, 3F). The asymmetry of the *θ* angle further increased during the looping stages (*d1*/*d2* ≥ 2) until reaching a 20º difference (Fig. 5B). In contrast, the *φ* angle left-right asymmetry was variable but, on average, did not differ significantly from symmetry at any stage (Fig. 5B). Furthermore, we found no obvious correlations between the L-R differences in *θ* and *φ* angles (Fig.5D). Therefore, before heart looping is obvious, and starting at late cardiac crescent stages, left-right morphological asymmetry builds up in the inflow region, manifested in the different angles of insertion of the inflows. Related to the different angle of insertion of the inflows, we frequently observed that the insertion of the right inflow into the ventricular region forms an acute angle, whereas the similar position on the left side is characterized by a much smoother transition, often even accompanied by a small bulge connecting the ventricle to the inflow (Fig. 5E). Again, these features can be detected from the cardiac crescent stages, but their presence is variable, with a proportion of the specimens in which this left-right difference is not obvious (Fig. 5E). In hearts at the looping stage, these asymmetric connections seem to favour the typical rightward ventricular displacement during looping (see specimen 47 in Fig. 5E), as the right-side acute angle may act as a hinge, favour rotation towards the right side, while the bulge on the left side may oppose a similar rotation towards the left side.

**Figure 5.**
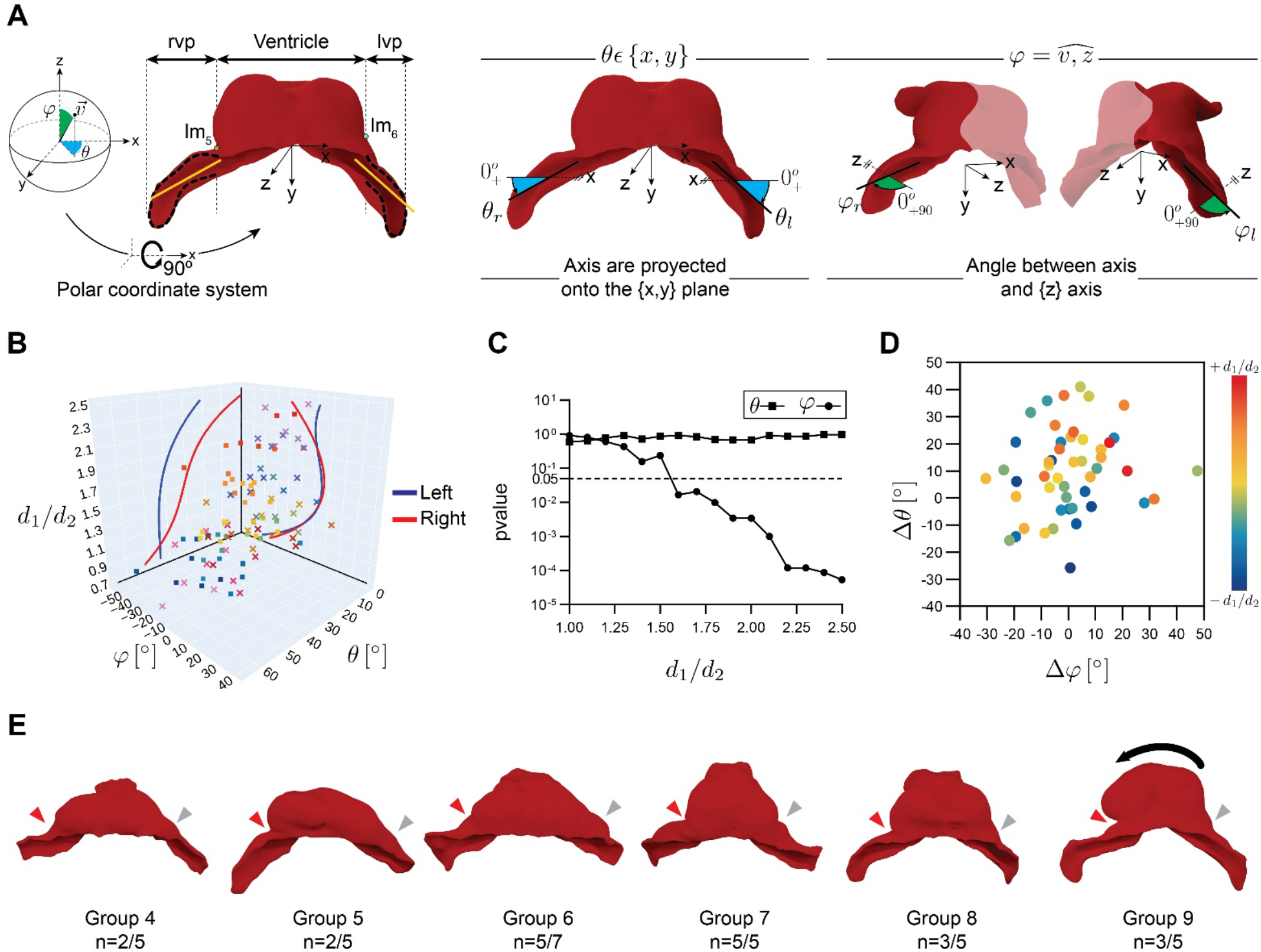
Onset of left-right morphological asymmetry before heart looping. (A) Schemes show the calculated principal directions of the inflow, labeled in orange. These directions are calculated as explained in Fig. S13. A polar coordinates system is then implemented for the full definition of the direction of the inflows by the magnitude of two angles, θ and the φ. (B) 3D plot showing the evolution of the θ and the φ angles in correlation with the parameter *d1*/*d2*. Lines the vertical planes of the graph show orthogonal projections of a polynomial curve adjusted to the trajectory of left (blue) and right (red) θ and φ angles. (C) Evolution of p-values for deviation from L-R symmetry of the θ and φ angles depending on the d1/d2 value. A mixed lineal model was applied to estimate the asymmetry between both sides at different stages (fixed effects) adjusting by the angle. (D) Scatter plots of the correlation values (Δθ, Δφ) for each specimen. (E) Specific hearts from different groups, highlighting the acute angle between the right inflow insertion and the ventricle (red arrow) and the smooth transition on the opposite side (grey arrow), often characterized by a small bulge. **n** indicates the number of specimens showing this feature over the total number of specimens in each staging group. Black arrow illustrates the direction of looping.

### A dynamic geometrical model of primitive heart tube formation and initial looping

To simulate temporal evolution through the stages characterized, we used the 10 MSG shapes generated plus specimen 51 (Fig. S7) and the method described in (Schmidt et al., 2020) to generate a smooth temporal transformation between stages (Fig. 6A). To achieve this, we propagated a common mesh connectivity from the first shape towards the latest one (Fig. 6A). Once the same mesh connectivity is set on all the MSGs, they were rigidly aligned to the first mesh (t1). In addition, a second alignment step was taken using the landmarks, in order to bring closer the same equivalent biological regions across stages. We then applied an interpolation technique (see material and methods) to estimate the positions of each vertex between stages, so that transitions between positions take place smoothly (Fig. 6C) and inserted 30 interpolated shapes between each MSG. This approach generated a dense temporal sequence of 3D meshes of heart tube morphogenesis (Movie 1; Dataset MSGs_4D). The model generated does not represent the evolution of a concrete specimen but a consensus most probable trajectory of the morphology of the forming heart tube, based on averaging the shapes of several hearts at each stage.

This approach did not allow us to include the models of the rest of tissues associated to the forming heart tube, because mismatches are generated at the limits between the MSGs of the different tissues. To generate a dynamic model that includes all the tissues, we then used the collection of 10 SGRs (Fig. 6B; Movies 2-7; Dataset SGRs_4D). Given that these are specific specimens, their matching to the surrounding tissues is perfect and the resulting dynamic model represents the shared boundary accurately, although occasional maladjustment artefacts can be observed. This model represents a possible evolution of the morphology of the heart tube and its associated tissues based on different specimens that represent consecutive stages of heart development. Examples of the interpolated tissues are provided in Fig. 6B.

**Figure 6.**
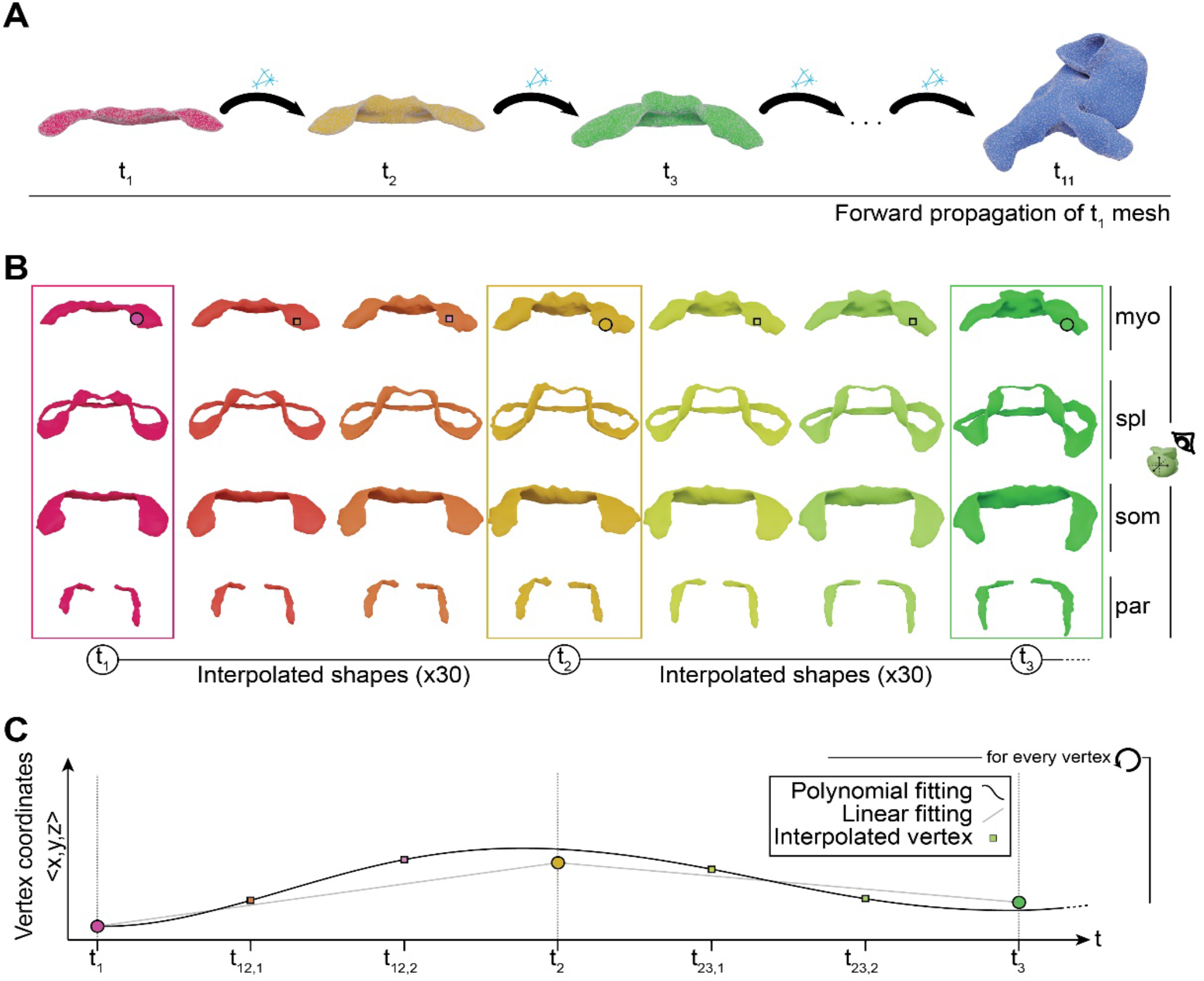
Dynamic meshes defining the full spatio-temporal domain of heart tube morphogenesis. (A) Panel shows the elaboration of a continuous sequence using the MSGs shapes. The process starts by mapping the mesh connectivity of the first timepoint to the next shape. This process is repeated until the connectivity of the first mesh is propagated up to the final mesh at t10. This process can also by applied when using the MSGs shapes. (B) The figure illustrates the concept of shape interpolation, applied to the three first timepoints of the SGRs shapes. Each row shows the result for the different tissues forming the pericardial cavity. The dots depicted on top of the myocardium shape illustrate the position of a vertex travelling through the shapes as morphogenesis progresses. (C) The plot illustrates the estimated trajectory of the vertex highlighted in panel B.

## DISCUSSION

Here we generated the first comprehensive dynamic and realistic representation of cardiac tissues during primitive heart tube formation in a mammalian embryo. To overcome the current limitations in live imaging of cardiac development at enough resolution and volume size, we opted for a pseudo-dynamic analysis based on the acquisition of large-volume, high-resolution 3D confocal images from a temporally dense collection of fixed specimens. We used a reporter line in which the membranes of all cells are labelled with high intensity fluorescence in different colours for the mesoderm and the rest of the tissues. The membrane labelling allowed a precise determination of the different cardiac tissues and their anatomical relationships with other tissues. Combining information of cell morphology, anatomical position and association to other tissue layers, we could determine the distribution of the main tissues involved in cardiogenesis with unprecedented detail. We reconstructed the differentiating myocardium, splanchnic mesoderm, somatic mesoderm, endoderm and endothelium. Beyond rendering the segmented tissues, we went further and generated finite-element models that recreate the geometry of each of the studied tissues with information on their inside-outside topology. In the process of generating a unified dynamic model, we faced the problem of morphological variability during early cardiogenesis, which represented limitations to staging and to the definition of “standard” models that would represent “normal” development. The high variability between specimens at similar developmental stages in fact means that it is not possible to determine a canonical shape trajectory for heart morphogenesis. As an alternative, we explored models that represent “most likely” geometries for each developmental stage. To achieve this, we first explored different approaches for morphometric staging that would not depend on subjective appreciation of specimens, which is the current state of the art in the field (see for example (Le Garrec et al., 2017; Tyser et al., 2021). Unsupervised clustering and assessment of parameters that measure the quality of classifications identified a simple relationship (d1/d2) as the best staging parameter. We think this staging method will be of utility in the precise definition of stages when comparing and 3D-mapping data between different laboratories. Furthermore, we identified simpler measurements accessible to all labs, like the height to width ratio, as very highly correlates of the d1/d2 parameter.

Ten staging groups were established according to the variation of d1/d2 and 9 of them contained 4 to 7 specimens, which allowed the statistical analysis of morphology. For this purpose, we used a recently developed tool (Schmidt et al., 2020) that finds dense pointwise correspondence between two surfaces of the same topology (genus). The use of this tool allowed the definition of mean shapes and the application of statistical analyses to each vertex, thereby generating maps of relative rates of local variability. We propose that this approach is of high value when describing mammalian organogenesis, since it simultaneously provides information about the “average” geometry of a developing organ and the regions that are more likely to vary from specimen to specimen. Primitive heart tube formation in amniotes is a highly complex process through which an initial monolayer of tissue of a crescent shape transforms into a tube oriented in a cranio-caudal direction and subsequently loops to achieve the definitive positions of the developing cardiac chambers. Failures in these early steps of development can lead to misalignment of the chambers and great vessels or to different types to left-right cardiac mispatterning. In particular, we identified strong variability at the dorsal borders of the myocardium. These dorsal borders, initially placed laterally, end up fusing at the dorsal midline, forming the dorsal mesocardium, which later disappears during looping. Important mechanical roles have been proposed for the mesocardium during looping (Le Garrec et al., 2017) and the mesocardial myocardium later localizes to the inner curvature of the looping heart, which plays essential roles in the alignment of the cardiac chambers and the great arteries (reviewed in (Gittenberger-de Groot et al., 2005)). The finding of high variability in this region therefore could be related to the onset of malformations associated to the mesocardial/inner curvature morphogenesis. Other regions of high variability were the inflows, which was expected, as these are highly dynamic and transient structures. Finally, apical regions of the future left ventricle show as well high variability, possibly associated with asynchrony of the ballooning process (Christoffels et al., 2000) between specimens.

Our approach also allowed us to study the left-right symmetry of the forming heart tube. Heart tube looping is considered as the first morphological evidence of organ left-right patterning in the embryo. In particular, cardiac left-right patterning defects are commonly associated with impairing congenital heart disease. Here we describe a new aspect of left-right asymmetry in the forming heart affecting the inflow tracts and preceding heart looping. The influence of the inflow region in heart looping has been proposed previously, mainly through the exertion of asymmetric forces from the left and right sides of the venous pole, provoked by differences in cytoskeletal contraction, cell migration and/or proliferation (Kidokoro et al., 2008; Ocana et al., 2017; Voronov et al., 2004). We therefore identified the first signs of morphological asymmetry in the mouse embryo localized to the inflow tracts of the forming heart. This asymmetry precedes looping and is likely to affect this process, as any active or reactive force exerted by/on structures oriented in different angles is predicted to produce asymmetric forces. We therefore propose that left-right bias in the orientation of the inflows depends on the general left-right patterning mechanisms of the embryo and influences the process of heart looping by imposing an architectural bias to the forces that shape it.

The dynamic models generated here represent highly detailed atlases of the different tissues involved in primitive cardiac tube formation. These atlases will be useful for the combination of relevant biological information produced from different experiments or labs into a unique spatial representation. This opens the door for establishing a common space where scientists can share their quantitative analyses on any quantitative aspect representable in the spatiotemporal domain, e.g.: gene expression patterns. This can be done by segmenting the complete or partial morphology of the heart, matching the stage with respect to the 3D+t atlas and map the data to the most similar shape of the atlas. Then, this mapping can be used to transfer values dwelling in either the reconstruction (scalar or vectorial values) or the original raw data (intensity values of a specific marker). Furthermore, the atlas generated is embedded in a statistical framework that accounts for natural variability, which can be used as a reference for analysis of mutant embryos. Our methodology for both global and local shape analysis, will allow to calculate differences either at global scale or affecting specific region between mutant and control specimens. This can be extended not only to the morphological level, but also for any quantitative magnitude expressed within the spatial domain of interest.

Understanding morphogenesis is one of the most active areas of research in developmental biology and here we generated tools that will help to understand mammalian cardiogenesis. Deep understanding of morphogenesis involves the identification of the sources and the temporal and spatial distribution of forces that shape developing tissues. Predictive modelling is ultimately required to fully acquire this knowledge (Sharpe, 2017); however, it critically relies on the access to realistic dynamic models of the geometry of the tissues/organs under study (Kawahira et al., 2020). The realistic mesh model of dynamic heart development we provide here will be useful for the development of finite-element models that explain the forces that drive cardiac tube formation, one of the most complex morphogenetic processes in the mammalian embryo, which underlies the high incidence of cardiac malformation in newborns.

## MATERIALS AND METHODS

### Animal model

Mouse alleles used in the manuscript are listed including bibliographic references and allele identities at the ‘Mouse Genome Informatics’ data base (MGI, http://www.informatics.jax.org/). *Mesp1*^*cre*^, MGI:2176467 (Saga et al., 1999), *Rosa26R*^*mTmG*^, MGI:3716464 (Muzumdar et al., 2007), *ROSA26R*^*Tomato*^, MGI:6260212 (Madisen et al., 2015) and C57BL/6 (Charles River). All animal procedures were approved by the CNIC Animal Experimentation Ethics Committee, by the Community of Madrid (Ref. PROEX 144.1/21) and conformed to EU Directive 2010/63EU and Recommendation 2007/526/EC regarding the protection of animals used for experimental and other scientific purposes, enforced in Spanish law under Real Decreto 1201/2005.

### Embryo preparation

Embryo collection was carried out at nights from 00:00 to 08:00 (E8-E8.33). Pregnant mice were euthanized by CO2. Individual embryos were isolated under a binocular from the uterus and the extra-embryonic membranes at room temperature. Embryos were then immersed in 0.1M phosphate buffered saline (PBS) and transferred to 2% paraformaldehyde fixative (PFA) overnight at 4ºC. After washing the embryos three times with fresh PBS to remove fixation medium, embryos were permeabilized in PBST (PBS containing 0.1% Triton X-100) for 30 minutes and washed in fresh PBS afterwards. Then embryos were incubated overnight at 4ºC with DAPI (1:500). After removing DAPI using fresh PBS, embryos were embedded in low melting point agarose dissolved in Milli-Q water at approximately 35ºC in 6-well plates. Embryos were kept in the desired orientation (Fig. 1A) using pipette tips, until agarose was solid enough. Then, the well plate was left for 30 minutes at RT, protected from light, until agarose solidifies completely. Then, the block of agarose was extracted from the well and put in the dissection microscope. Using a sharp knife, a small cube was created (0.5cm x 0.5cm x 0.5cm) containing the embryo. The cube was then immersed in CUBIC I during 1 day at 4ºC. Then CUBIC I was renewed and left incubating for 2 days at 4ºC.

### Embryo imaging

Embedded embryos were imaged in an inverted microscope through an 80µm-thick cover glass placed on a perforated culture plate, to minimize optical pathway length (Fig. 1A). Then, a pipette was used to aspirate excess CUBIC so that the face of the agarose cube is in direct contact with the glass, thereby reducing optical pathway length. Given the high sensitivity of CUBIC I to water concentration, a saturated atmosphere was created to prevent evaporation. This was done by placing a couple of 1.5 ml plastic tubes filled with water in the culture dish and covering the culture dish.

Confocal images were obtained using a SP8 Leica confocal microscope and a HC PL Apo CS2 20 0.75 NA multi-inmersion lens corrected to glycerol immersion media was used. Scanning settings: xyz mode, bidirectional, 400Hz. Pinhole: 56.7µm. Zoom: 1.15. Laser lines used: diode 405nm (5% power, DAPI signal), 488nm (power 15%, mGFP signal), 561nm (20%, Tomato signal). Detectors used: PMT (424nm-474nm), HyD (496nm-529nm, gain 100% and 583nm-667nm, gain 170%).

When the size of the embryo could not be covered by the configuration described, a tile-scan was performed, and tiles were stitched using Leica LAS X 3.5.2.18963 software. Raw images were deconvolved with “Huygens Professional version 19.10”, to compensate for the blurring and noise inherent to the optical system.

Segmentations were performed in a 1024×1024 resized version of the raw images, which were previously rotated in 3D to orient them with respect to the embryo axis (Schindelin et al., 2012).

As the acquisition is performed in the confocal microscope, left and right side are inverted. After segmentation images are converted to surfaces, these are flipped with respect to the y axis of the image, so that all surface meshes are oriented as indicated in the system of reference shown in Fig. 1A.

### Embryo collection

We have imaged a total of 52 specimens. Within these data, there are some exceptions in which we have not been able to obtain all the tissues and shapes of interest (Fig. 1A). In the following, we list the name of the embryo, its corresponding staging group and what makes the dataset exceptional.

- E21, group 5: we have not been able to reconstruct the somatic or paraxial mesoderm of this embryo, as this area fractured during embryo handling.
- E22, group 5: we have not been able to image the most dorsal part of the foregut endoderm, neither the aortic lumen in that region.
- E26, group 6: we have not been able to fully reconstruct the foregut endoderm, as this embryo presented a small break in the ventral side of the foregut endoderm.
- E37, group 8: we have only been able to segment the morphology of the myocardium and the endocardial lumen.
- E41, group 8: we have not been able to segment the splanchnic and somatic mesoderm because both were broken at some points. We have been able to accurately segment the myocardium.
- E51, outlier: we have not been able to segment the foregut endoderm and the aorta. This is because in this very advanced specimen, the cardiogenic area extends in the microscopic z axis longer that we have been able to reach with the imaging protocol.

### Exceptions on morphological events timing

- E29: The myocardium of this embryo is closed prematurely.
- E32: This heart is apparently looping prematurely, however looping is apparently reversed.
- E35: This heart is apparently looping prematurely.
- E42, E46: These hearts are dorsally closed, while overall morphology reflects an earlier stage.
- E44, E49: Despite looping is advanced in these specimens, dorsal closure is far from being complete.

### Surface map computation

We computed surface maps using the method described in (Schmidt et al., 2020). This method takes two topologically equivalent surface meshes and a sparse set of corresponding points (landmarks) as input. As output, it produces a dense map that determines for every point on one surface a corresponding point on the other surface, and vice versa. In particular, these maps are continuous and bijective, meaning that problematic artefacts such as tearing or fold-overs (multiple points mapped to the same location) cannot occur. The algorithm minimizes intrinsic mapping distortion, i.e., it reduces the amount of stretch that each piece of one surface undergoes when being mapped to the other surface. Minimizing mapping distortion often leads to alignment of geometrically similar surface regions (Schmidt et al., 2019). We initialized each map via the layout embedding method presented in (Born et al., 2021).

### Shape averaging

Computing a mean shape within each group requires surface correspondence, i.e., which point on one shape to average with which points on the other shapes. To this end, we selected one reference shape per group, whose triangulation serves for the mean shape (Fig. 4A’). We then computed surface maps (Schmidt et al., 2020) from the reference to all other shapes of the group. We used each map to transfer the vertices of the reference mesh to the respective target surface, establishing a vertex-to-point correspondence. Using this correspondence, we rigidly aligned all shapes of the group to the reference shape to compensate rotational and translational misalignments. We then computed the average 3D position per vertex of the reference mesh, i.e., the mean of the reference vertex itself and its corresponding points on the other rigidly-aligned shapes (Fig. 4A’). As a measure of local shape variability within the group, we report the standard deviation per set of corresponding vertices, given by the square root of the largest eigenvalue of the covariance matrix (Fig. 4A’).

### Sequential mapping

Given an ordered sequence of shapes, we established correspondence across this sequence by computing a surface map between each consecutive pair of shapes and then successively composing maps.

As all shapes exhibit different triangulations, it is necessary to establish a common reference triangulation for the entire sequence. We initially chose the triangulation of the first shape for this purpose and successively transferred its vertices to all surfaces of the sequence (Fig. 6). The result is a single triangle mesh connectivity where each vertex is equipped with a set of positions, one for each shape of the sequence. These act as the keyframes of a continuous description of the morphological process. Again, we rigidly aligned all shapes of the sequence in 3D to compensate rotational and translational misalignments between keyframes.

### Compatible remeshing

The initial reference triangulation (transferred from the first shape to the rest of the sequence) does not provide a high-quality surface approximation at every stage of the morphological process. To this end, we employed a remeshing technique based on (Botsch and Kobbelt, 2004) to obtain a common triangulation that is well suited for all keyframe shapes. We computed a target edge length field on each shape of the sequence (Dunyach et al., 2013). We then used sequential surface maps to combine all fields via a vertex-wise minimum. That is, if at least one shape in the sequence requires a high mesh resolution in some area, this demand will be respected in the common triangulation. We then iteratively improved the common triangulation by a series of edge splits, collapses, flips and tangential Laplacian smoothing (Botsch and Kobbelt, 2004), while simultaneously considering the reference mesh embedded on all shapes of the sequence (Yang et al., 2020).

### Shape interpolation

To approximate a continuous morphological evolution, we uniformly sampled 30 new timepoints between each pair of consecutive shapes in the sequence. We initially determined the intermediate shapes via linear interpolation and then applied a polynomial smoothing filter to elude sharp transitions (Luo et al., 2005). We provide a collection of 11 timepoints plus 30 timepoints between each frame, which represents a total of 311 shapes.

### Calculation of the dorsal closure gap

The procedure uses the landmarks 9 and 7, and 8 and 10 to define the edges of the dorsal lips of the myocardium (labelled as yellow in Fig. S11). The dorsal gap, *l*, measures the length of the shortest line that joins both lips. The calculation of *l* is achieved by the subdivision of the edge of the lips, generating N points in the left side and K points in the right side. Then, for every point *j* (left lip) to any point on the right lip, the minimum distance is calculated, d(p_j_, q). Finally, the minimum of the calculated distances is taken as the dorsal gap.

### Surface map computation between shapes of different topology

The dorsally closed heart and the splanchnic mesoderm represent cases of torus topology (genus 1), while all others have sphere topology (genus 0) This represents a compatibility problem when computing surface maps across the entire sequence. Therefore, we have developed a methodology to turn these shapes into sphere topology (genus 0) surfaces, by cutting the closed surfaces in a specific region. Regarding the dorsally closed heart, the border between the two myocardial lips is considered as the cut trajectory (Fig. 4B, yellow dotted-line and arrow). In the case of the splanchnic, a cut is made along the AP axis, in the medial domain, anterior to the outflow tract (Fig. 1G, black dotted-line and arrow). For the latter, this procedure was done to all the splanchnic shapes of the collection since they all present 1-genus topology.

We processed each cut by individually triangulating the two emerging holes, yielding a closed surface of genus 0. To fuse any visible gap, we slightly extruded both newly triangulated areas (producing a subtle self-intersection of the geometry), and finally remeshed the affected regions.

### Mid-surface extraction

The meshes were first isotropically remeshed (Cignoni et al., 2008) to 2µm edge length, and smoothed (Taubin, 1995). Then, the skeleton of the mesh was derived using (Yoshizawa et al., 2003) and smoothed to eliminate spikes and/or high-frequency details. The former step was repeated until the algorithm creates a mono-layer mesh. Then, the resulting mesh was resampled using a Poisson disc sampling and reconstructed using the Ball pivoting algorithm (Cignoni et al., 2008). After this, the mesh was smoothed and post-processed to eliminate imperfections (holes, non-manifold edges, non-manifold vertices, etc.). The final state of the mid-surface comes given by a last isotropic explicit remeshing (2µm).

### Definition of the inflow insertion angles

Fig. S13 illustrates and explains the steps taken to generate a system of reference inherent to every myocardium shape (mSOR, myocardium system of reference), as well as the strategy to estimate the principal direction of the inflows. The insertion angles are the result of calculating the angle of the principal direction with respect to the system of reference (Fig. 5A). First, a polar system of reference (PSoR) is defined, together with the angles *θ* and *φ*, which define the unit vector, *v*. Then, the PSoR is translated to the mSOR. The *θ* angle (Fig. 5A, middle) is defined within the plane formed by the {x,y} plane, whose normal is approximately equal to the cranio-caudal axis of the embryo. In order to obtain comparable values of *θ* from both inflows, the origin of the angles is defined differently in each side, as denoted the 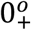 terms. The *φ* angle (Fig. 5A, right) is defined as the angle through which an inflow pivots with respect to the dorsal-ventral axis. This angle is calculated as the angle formed between the principal direction of the inflow and the z axis. As this angle is defined, it would oscillate around a value 90º. So, and offset of −90º is applied, as denoted by the 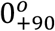.

### Definition of landmarks for map surface computation and shape registration

A set of landmarks has been chosen for every individual tissue. These landmarks have been used for the initialization of the surface map computation and for the registration of the MSGs and SGRs shapes. The landmarks are points consistent and clearly recognizable through the collection. An additional set of landmarks has been set by uniformly subdividing the curves formed by the mid-surface boundaries. This extra set of points is used in conjunction with the original landmarks, for a better performance of the alignment steps.

#### Landmarks in the myocardium (Fig. S11C)

Landmarks lm1, lm2, lm3 and lm4 have been set at the lateral ends of the inflows, at the places where the cardiac mesoderm is separated from the splanchnic. lm5 and lm6 are located at the point where we see the inflows separate from the ventricle. lm7, lm8, lm9 and lm10 are located at the border of the dorsal closure lips. lm11 is located at the most cranial point of the intersection between the mid-sagittal plane and the outflow. lm12 is located at the intersection between the mid-sagittal plane and the intersection between the myocardium and the juxta-cardiac field. The curves extracted from the edge of the mid-surface and using some landmarks are: curve1, goes from lm3 to lm4; curve2, goes from lm1 to lm2; curve3, goes from lm1 to lm3 and lm2 to lm4 on the left and right side respectively.

#### Landmarks in the splanchnic (Fig. S11D)

Landmarks lm3, lm4 and lm12 are the same as in the nearest myocardial landmark, mapped to the closest vertex of the splanchnic shape. lm13 is defined at the intersection of the splanchnic with the somatic mesoderm, at the point where the mid-sagittal plane intersects. lm15 is defined at the cranial-most point of the intersection between the mid-sagittal plane and the splanchnic. Landmark lm14 is obtained by calculating the midpoint of the geodesic curve that joins lm12 and lm13. Lm16 and lm17 are located at the bilateral most posterior caudal points of the splanchnic, at the right and left side respectively. lm24 and lm25 result from the calculation of the midpoint of the geodesic curve that joins lm16 and lm4 or lm17 and lm3, respectively. The curves extracted from the edge of the mid-surface and using some landmarks are: curve4, labels the border between the splanchnic and somatic mesoderm; curve5, is a bilateral curve joining lm24, lm15 and lm25 (left); curve6, joins lm24 and lm25 running equidistant between curve2 and curve4; curve7 is similar to curve 5 but running equidistant between curve 1 and curve 5, so that it runs through the mid-line of the second heart field.

#### Landmarks in the somatic mesoderm (Fig. S11E)

Landmarks lm13 and lm15 are located at the intersection of the outer edge of the somatic mesoderm with the mid-sagittal plane. Landmark lm26 is obtained by calculating the midpoint of the geodesic curve that joins lm13 and lm15. Lm18 and lm19 are the most caudal points of the somatic tissue. The curves extracted from the edge of the mid-surface and using some landmarks are: curve8, labels the border between the somatic and the splanchnic mesoderm; curve9, starts at lm18 and ends at lm19, it runs over the border between somatic and the paraxial mesoderm; curve 10, starts at lm18 and ends at lm19 and is defined as the equidistant line between curve 8 and curve 9.

#### Landmarks in the head paraxial mesoderm (Fig. S11F)

lm20 and lm21 are defined at the caudal-most points of the right and left domains of the paraxial mesoderm, respectively. Lm22 and lm23 are defined at the cranial-most points of the right and left domain of the paraxial domain, respectively. The curves extracted from the edge of the mid-surface and using some landmarks are: curve11, is a bilateral curve running between the paraxial and somatic mesoderm; curve12, is a bilateral curve running between the paraxial and the splanchnic mesoderm; curve13, runs equidistant between curve 11 and curve 12.

### Estimation of the mid-sagittal plane

Estimating the mid-sagittal plane (Fig. S11B) is necessary for the determination of some landmarks. The raw data is analyzed, and we manually assign points in regions included in the mid-sagittal plane of the embryo. When this number of points is larger than 3, the midplane is estimated by calculating the origin and normal of the plane that best fit those points. As a result, all the tissues can be assigned with respect to this plane.

### Renderings

All the renderings have been processed in blender 2.92 (https://www.blender.org/).

### Movies

**Movie 1**. The movie represents the continuous model of the myocardium, elaborated with the mean myocardium shapes.

**Movie 2**. The movie represents the continuous model of the myocardium, elaborated with the representative specimen of each staging group.

**Movie 3**. The movie represents the continuous model of the splanchnic mesoderm, elaborated with the representative specimen of each staging group.

**Movie 4**. The movie represents the continuous model of the somatic mesoderm, elaborated with the representative specimen of each staging group.

**Movie 5**. The movie represents the continuous model of the paraxial mesoderm, elaborated with the representative specimen of each staging group.

**Movie 6**. The movie represents the continuous model of the myocardium (red), in combination with the splanchnic mesoderm (blue), elaborated with the representative specimen of each staging group.

**Movie 7**. The movie represents the continuous model of all the tissues forming the anterior celomic cavity, elaborated with the representative specimen of each staging groups. Myocardium, red; splanchnic mesoderm, blue; somatic mesoderm, green; paraxial mesoderm, purple.

## Acknowledgements

We thank members of the Torres laboratory for fruitful discussions and advice on this work. We thank the CNIC Microscopy & Dynamic Imaging Unit, supported by FEDER, “Una manera de hacer Europa”. We are grateful to Fátima Sánchez-Cabo for helpful advice on the statistics. Isaac Esteban acknowledges attendance to the Computational Image Analysis (CIAN) course at the MBL, Woods Hole, Massachusetts (funded by NIH grant R25 GM103792)”.

## Author Contributions

Conceptualization: I.E., M.T.; Methodology: S.T., I.E., P.S.; Formal analysis: I.E.; Investigation: S.T.; Writing - original draft: I.E., M.T.; Writing - review & editing: I.E., P.S., L.K., M.T.; Supervision: L.K., M.T.; Funding acquisition: M.T.

## Data availability

Datasets with segmentation images, 3D shapes and 3D+”t” models are available at: https://data.mendeley.com/datasets/t828xhg66k/draft?a=1507233b-9c38-48ba-bdad-bffb6f4731b0

---

**Supplementary Figure 1.**
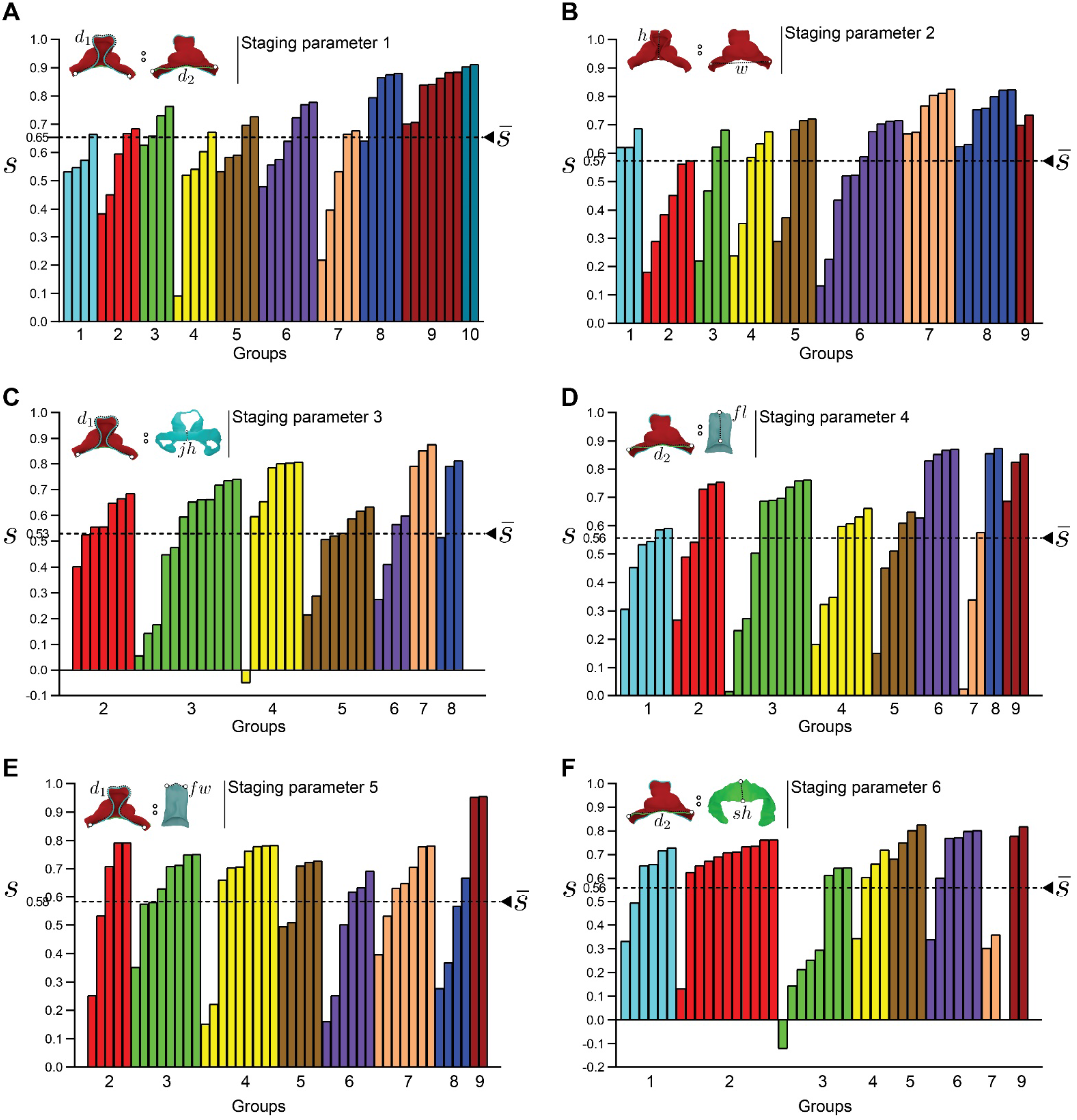
Silhouette coefficient plot for the different staging parameters explored, classified using k-means and number of groups equal to 10. (A-F) Bar graphs showing the silhouette coefficient value (*s*) for each embryo in the collection classified into 10 groups, according to the different staging parameters. The graphs also show the average value 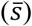, which is maximal for parameter 1. Graphs are color-coded according to the table shown in Fig. 3D. Schemes above the graphs show a representation of staging parameter used.

**Supplementary Figure 2.**
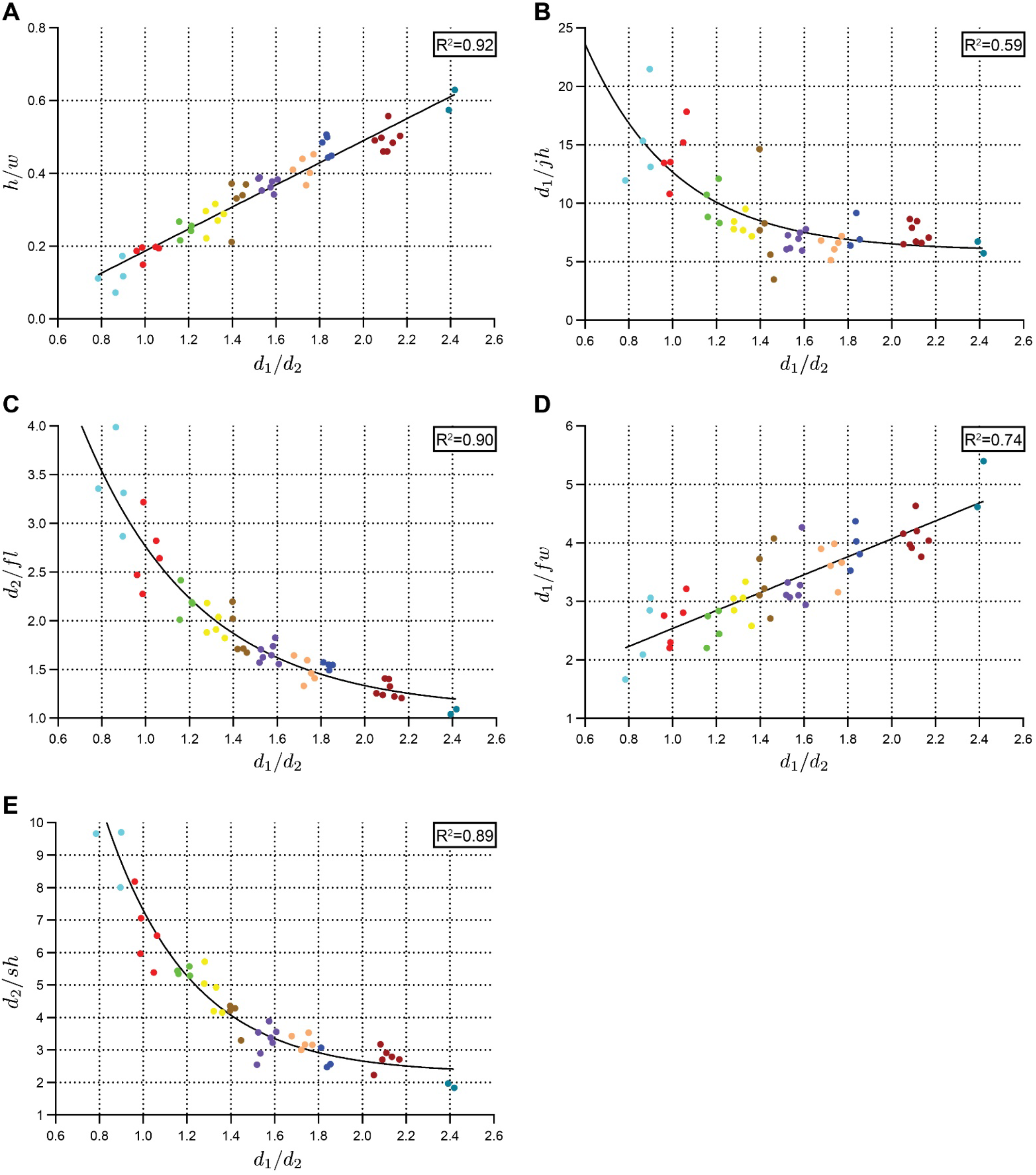
Correlations between staging parameters unveil tight relationships between the evolution of different morphometric aspects during heart development. (A, B, C, D, E) Plots show the correlation of parameters 2-6 (Y axes) to parameter 1 (d1/d2, X-axis).

**Supplementary Figure 3.**
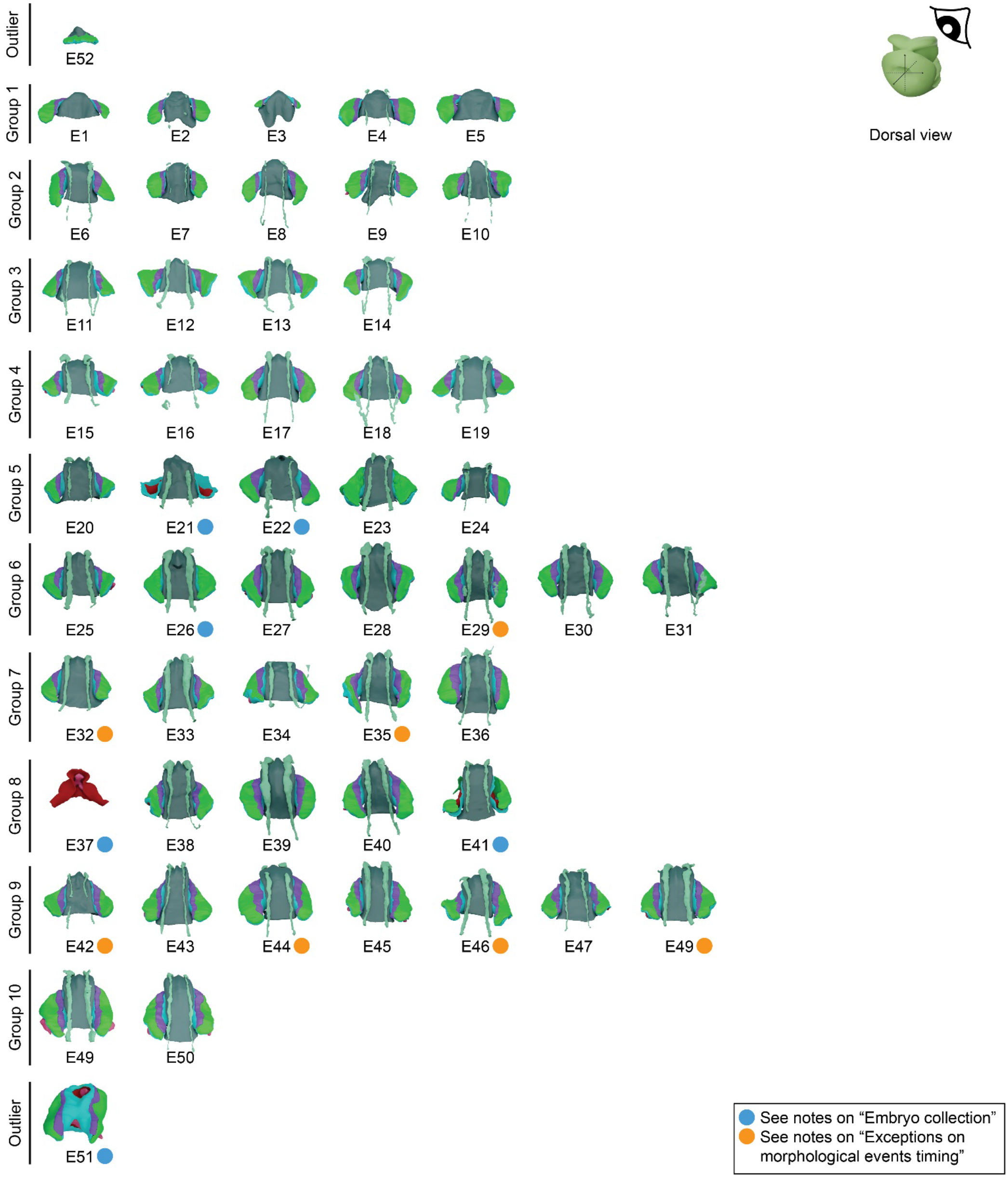
Embryo collection classified by staging groups, showing a dorsal view of all tissues. Dorsal view of all the specimens in the collection with representation of the processed surfaces of the complete set of tissues and shapes as described in Fig. 1G. The specimens at the very top and bottom are outliers, left out of the stage classification.

**Supplementary Figure 4.**
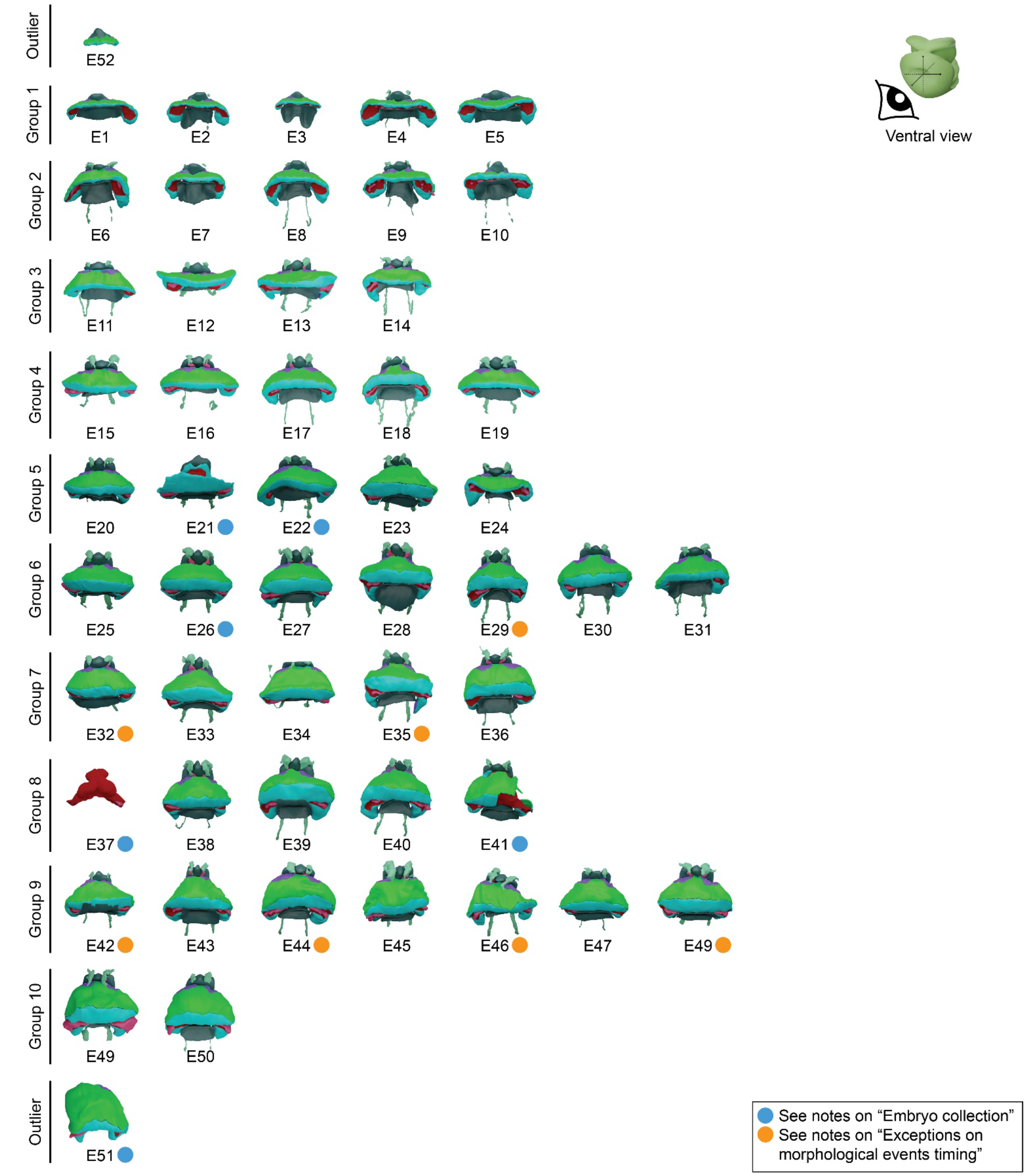
Embryo collection classified by staging groups, showing a ventral view of all tissues. Ventral view of all the specimens in the collection with representation of the processed surfaces of the complete set of tissues and shapes as described in Fig. 1G. The specimens at the very top and bottom are outliers, left out of the stage classification.

**Supplementary Figure 5.**
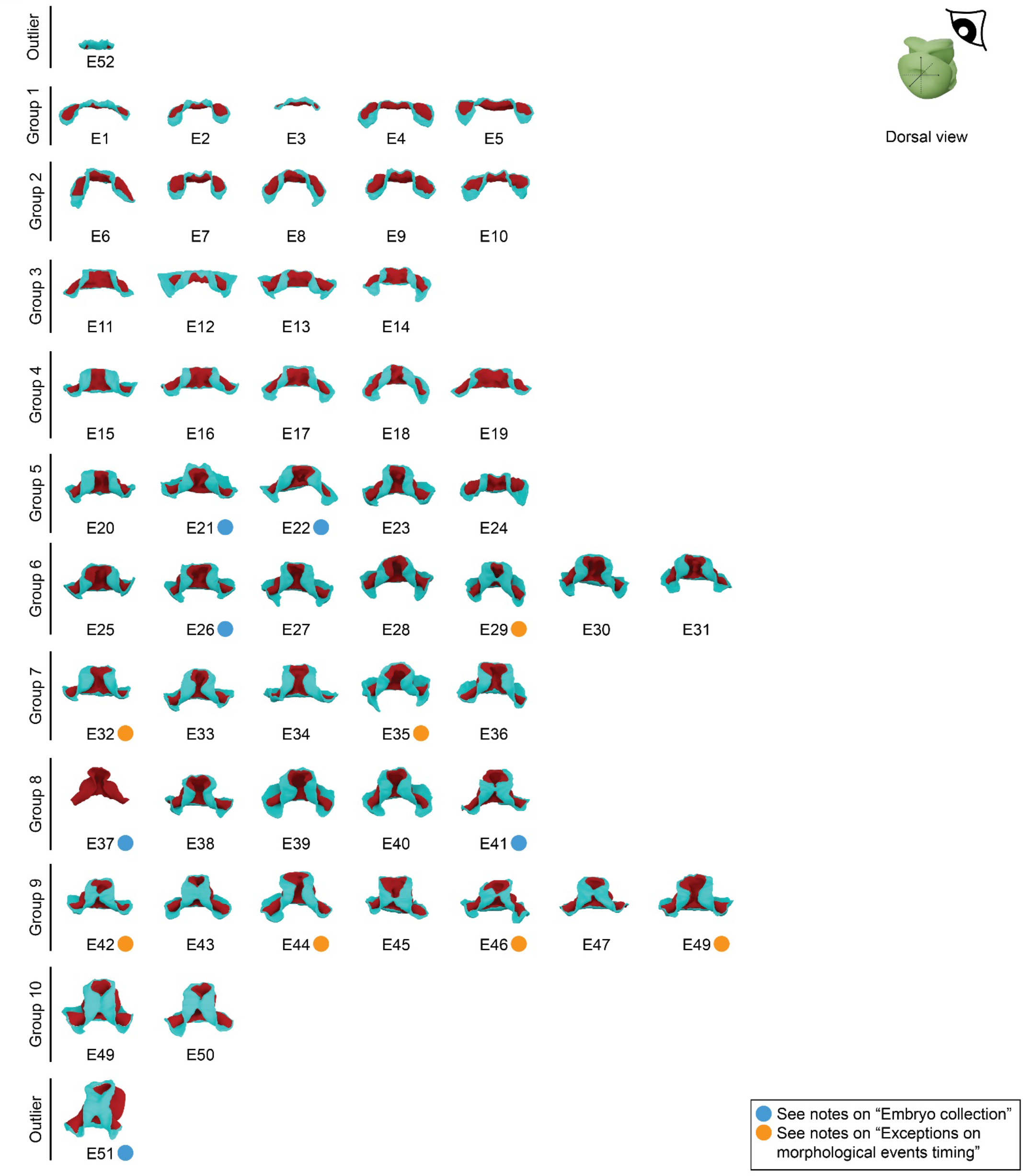
Embryo collection classified by staging groups, showing a dorsal view of splanchnic mesoderm and myocardium. Dorsal view of all the specimens in the collection with representation of the processed surfaces of the splanchnic mesoderm and myocardium. The specimens at the very top and bottom are outliers, left out of the classification.

**Supplementary Figure 6.**
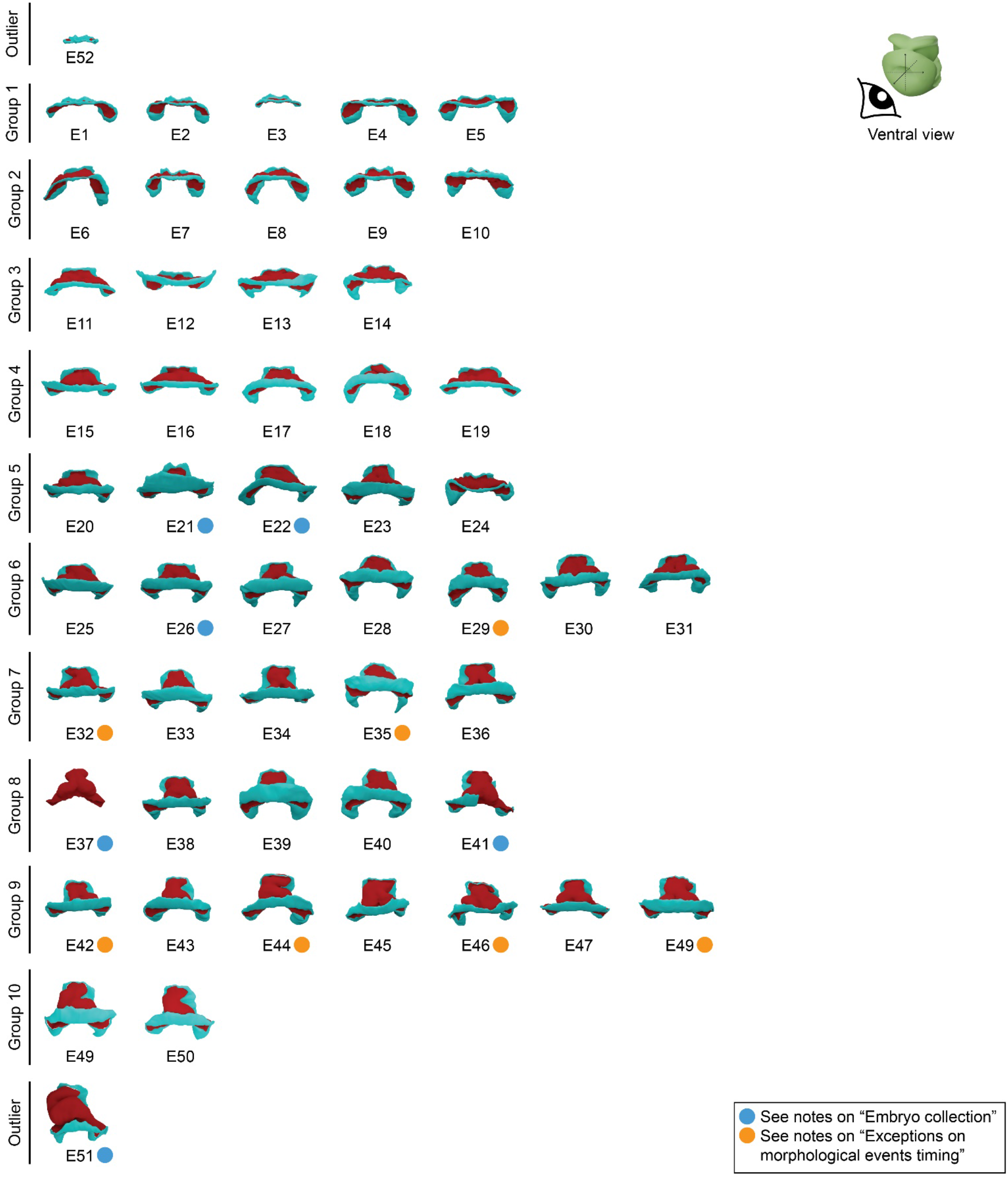
Embryo collection classified by staging groups, showing a ventral view of splanchnic mesoderm and myocardium. Ventral view of all the specimens in the collection with representation of the processed surfaces of the splanchnic mesoderm and myocardium. The specimens at the very top and bottom are outliers, left out of the classification.

**Supplementary Figure 7.**
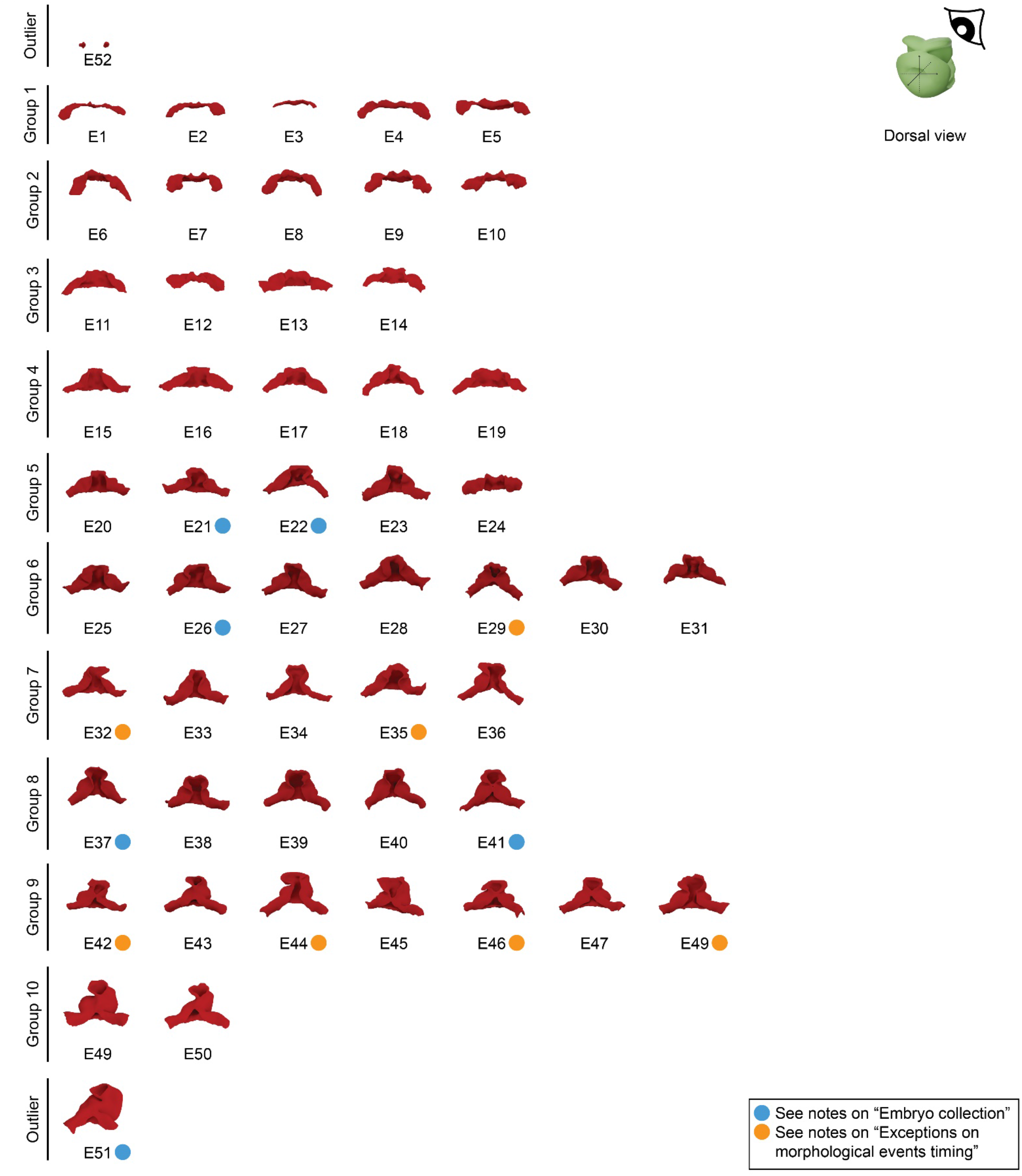
Embryo collection classified by staging groups, showing a dorsal view of the myocardium. Dorsal view of all the specimens in the collection with representation of the processed surfaces of the differentiated myocardium. The specimens at the very top and bottom are outliers, left out of the classification.

**Supplementary Figure 8.**
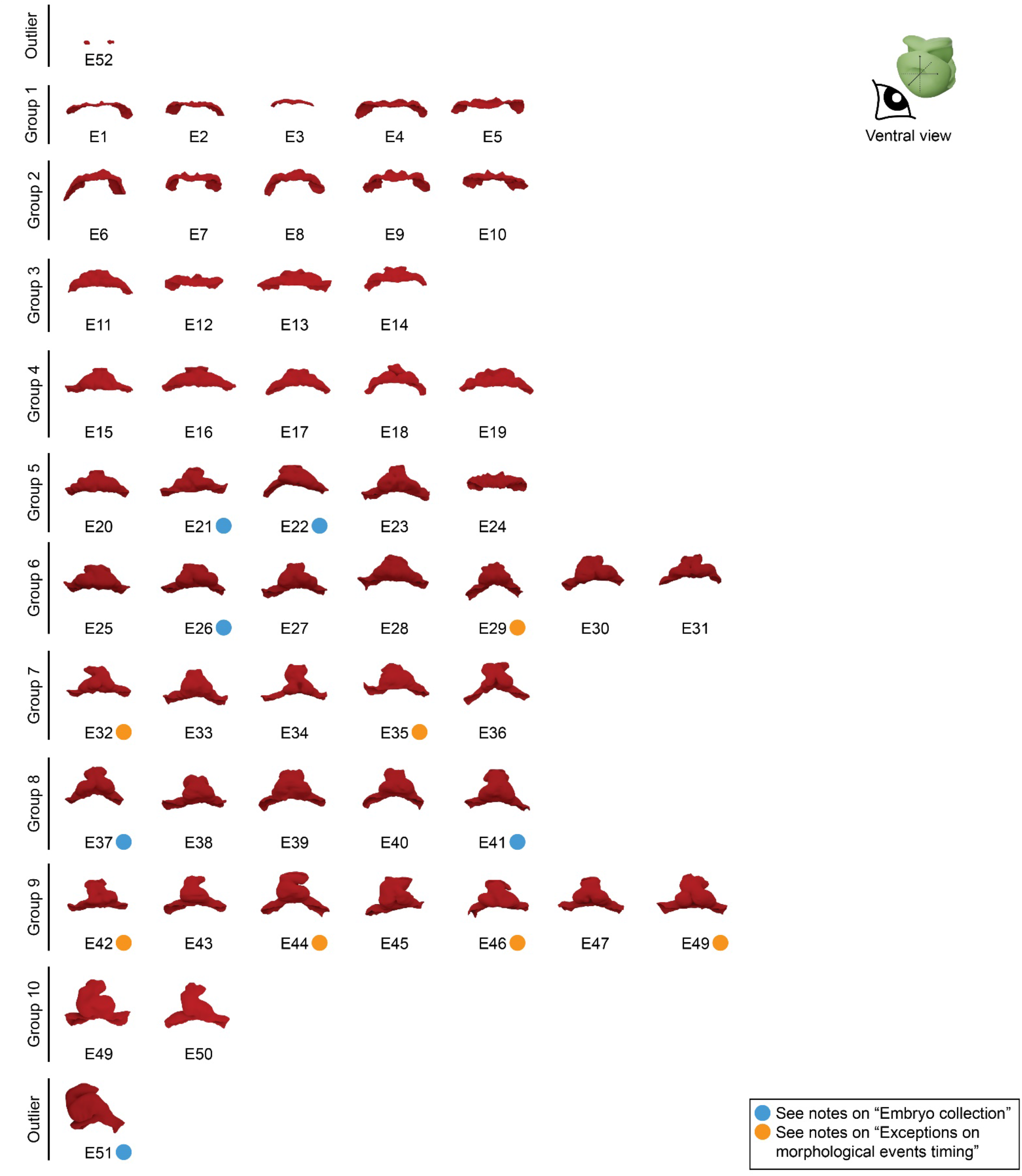
Embryo collection classified by staging groups, showing a ventral view of the myocardium. Ventral view of all the specimens in the collection with representation of the processed surfaces of the differentiated myocardium. The specimens at the very top and bottom are outliers, left out of the classification.

**Supplementary Figure 9.**
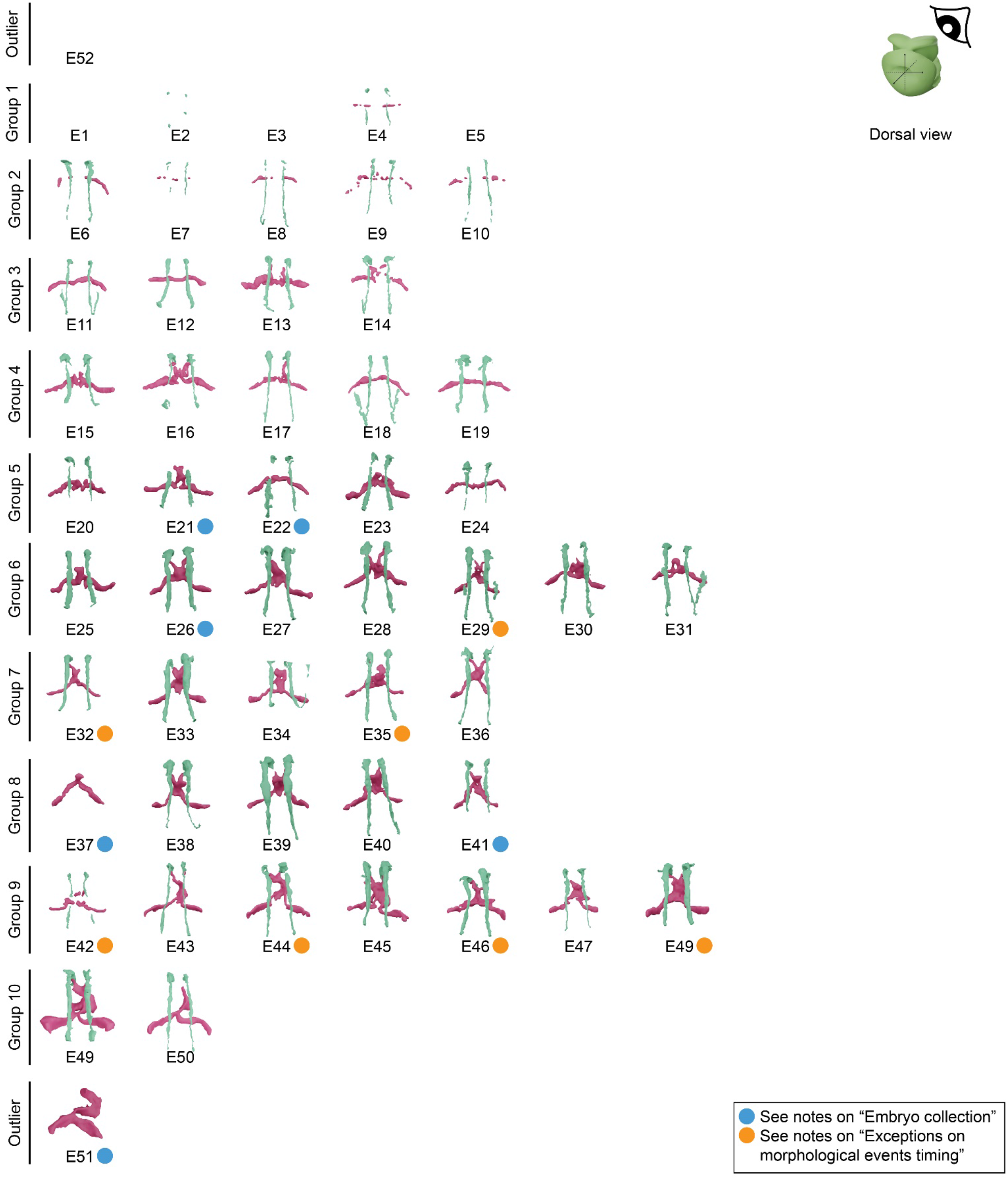
Embryo collection classified by staging groups, showing a dorsal view of the circulatory system. Dorsal view of all the specimens in the collection with representation of the processed surfaces of the circulatory system, split in two parts: endocardial lumen and aortic lumen. The specimens at the very top and bottom are outliers, left out of the classification.

**Supplementary Figure 10.**
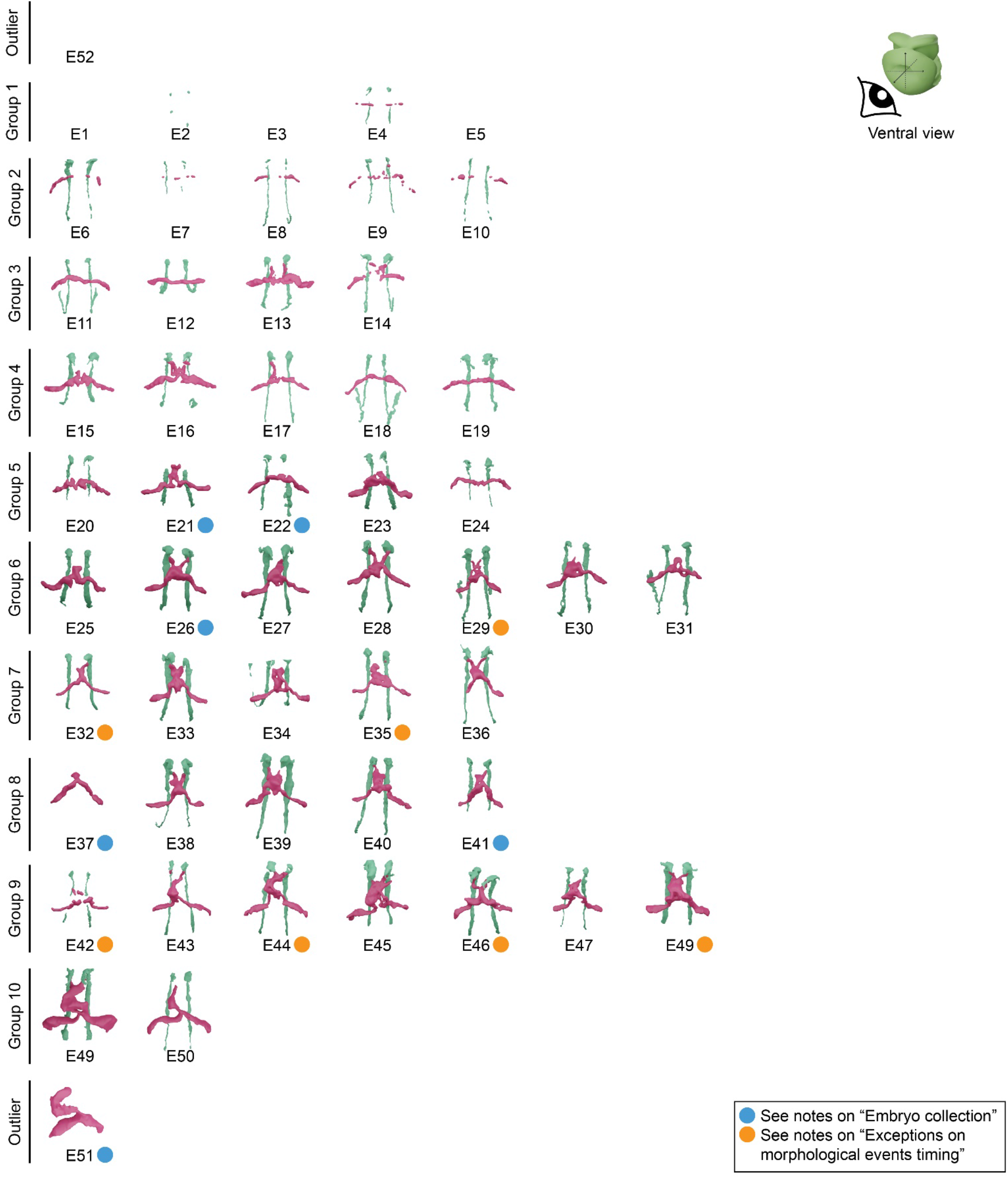
Embryo collection classified by staging groups, showing a ventral view of the circulatory system. Ventral view of all the specimens in the collection with representation of the processed surfaces of the circulatory system, split in two parts: endocardial lumen and aortic lumen. The specimens at the very top and bottom are outliers, left out of the classification.

**Supplementary Figure 11.**
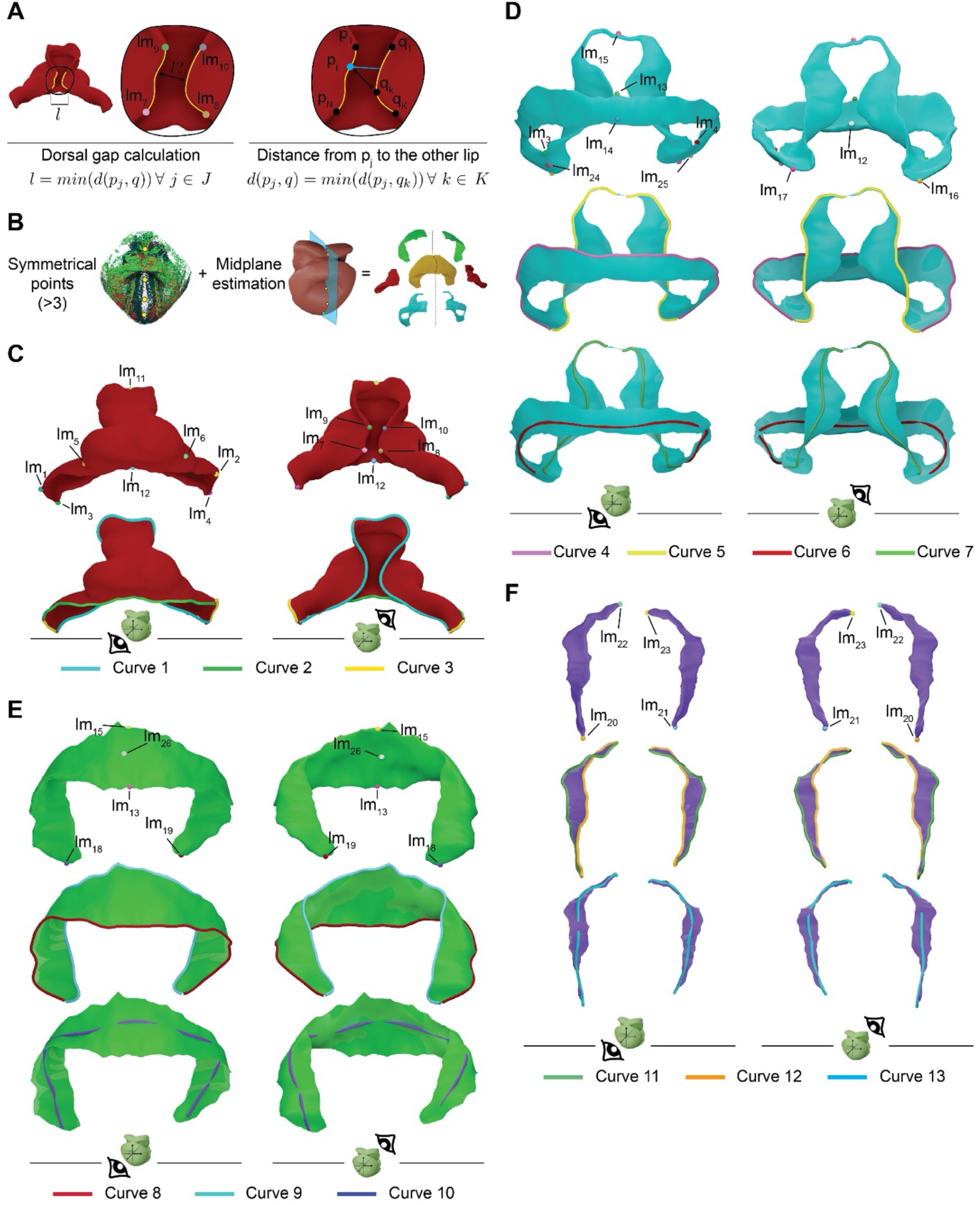
Measurement of dorsal closure and definition of landmarks and curves for surface map computation and shape registration. (A) Schematic representation of the procedure for estimating the dorsal closure gap. (B) This panel shows the process for estimating the mid-sagittal plane midplane of the embryo. The right part shows the tissues of the pericardial cavity cut in half trough the plane. (C) The panel shows the basic set of landmarks defined for the myocardium. (D) The panel shows the basic set of landmarks defined in the splanchnic mesoderm. (E) The panel shows the basic set of landmarks defined in the somatic mesoderm. (F) The panel shows the basic set of landmarks defined in the head paraxial mesoderm.

**Supplementary Figure 12.**
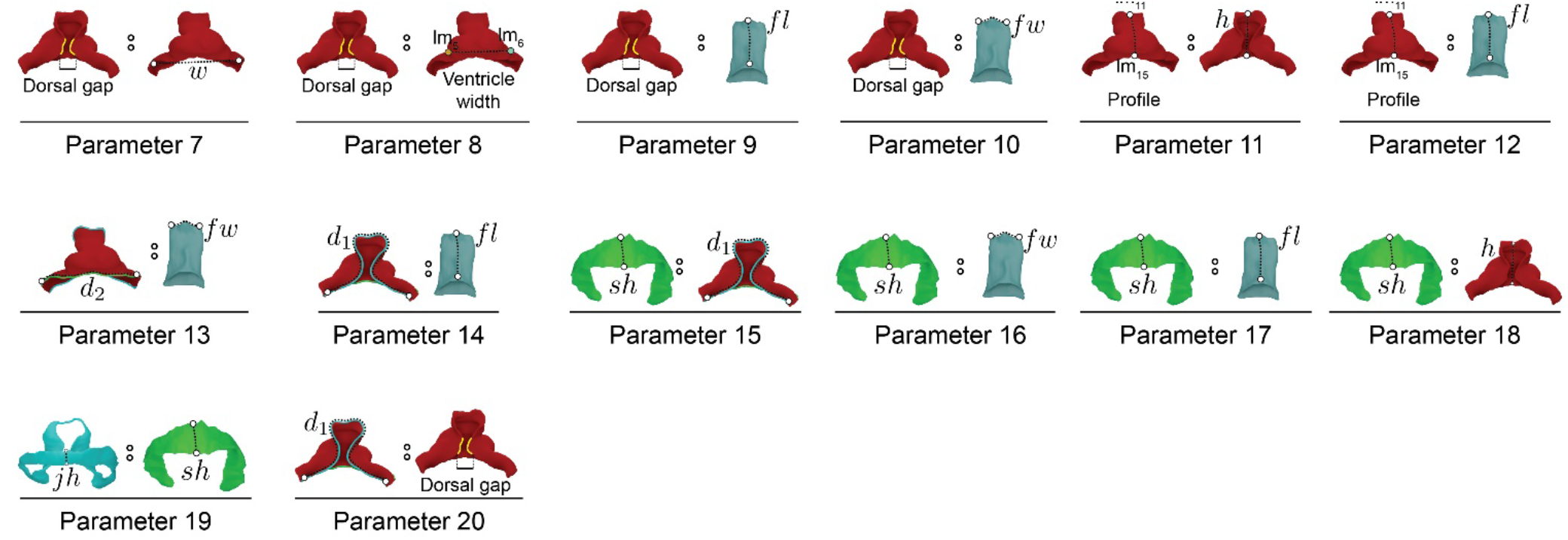
Additional parameters explored for staging. Schematic representation of additional parameters explored for the staging system.

**Supplementary Figure 13.**
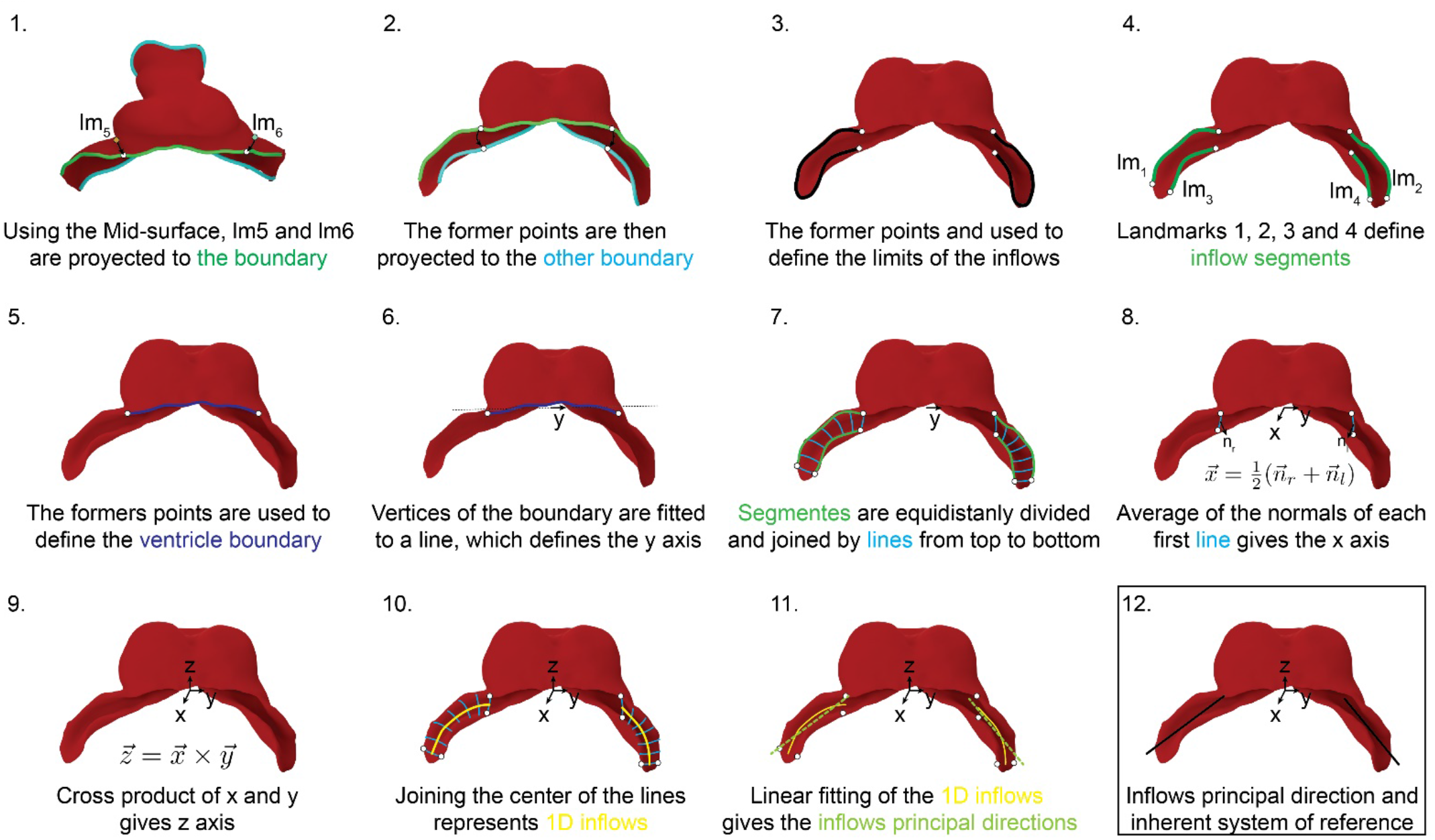
Proposed algorithm for systematic calculation of a myocardium inherent system of reference and calculation of inflow insertion direction. Steps followed for determining the principal direction of the inflows and a system of reference, using objective morphology.

## REFERENCES

Born, J., Schmidt, P. and Kobbelt, L. (2021). Layout Embedding via Combinatorial Optimization. Computer Graphics Forum 40, 277–290.

Botsch, M. and Kobbelt, L. (2004). A remeshing approach to multiresolution modeling. In Proceedings of the 2004 Eurographics/ACM SIGGRAPH symposium on Geometry processing, pp. 185–192. Nice, France: Association for Computing Machinery.

Christoffels, V. M., Habets, P. E., Franco, D., Campione, M., de Jong, F., Lamers, W. H., Bao, Z. Z., Palmer, S., Biben, C., Harvey, R. P., et al. (2000). Chamber formation and morphogenesis in the developing mammalian heart. Developmental biology 223, 266–278.

Cignoni, P., Callieri, M., Corsini, M., Dellepiane, M., Ganovelli, F. and Ranzuglia, G.(2008). MeshLab: an Open-Source Mesh Processing Tool.

de Bakker, B. S., de Jong, K. H., Hagoort, J., de Bree, K., Besselink, C. T., de Kanter, F. E., Veldhuis, T., Bais, B., Schildmeijer, R., Ruijter, J. M., et al. (2016). An interactive three-dimensional digital atlas and quantitative database of human development. Science 354.

de Boer, B. A., van den Berg, G., de Boer, P. A., Moorman, A. F. and Ruijter, J. M. (2012). Growth of the developing mouse heart: an interactive qualitative and quantitative 3D atlas. Developmental biology 368, 203–213.

Dunyach, M., Vanderhaeghe, D., Barthe, L. and Botsch, M. (2013). Adaptive Remeshing for Real-Time Mesh Deformation.

Faber, J. W., Hagoort, J., Moorman, A. F. M., Christoffels, V. M. and Jensen, B. (2021). Quantified growth of the human embryonic heart. Biology open 10.

Gittenberger-de Groot, A. C., Bartelings, M. M., Deruiter, M. C. and Poelmann, R. E. (2005). Basics of Cardiac Development for the Understanding of Congenital Heart Malformations. Pediatric Research 57, 169–176.

He, K., Gkioxari, G., Dollár, P. and Girshick, R. (2017). Mask R-CNN. In 2017 IEEE International Conference on Computer Vision (ICCV), pp. 2980–2988.

Ivanovitch, K., Temiño, S. and Torres, M. (2017). Live imaging of heart tube development in mouse reveals alternating phases of cardiac differentiation and morphogenesis. eLife 6.

Kawahira, N., Ohtsuka, D., Kida, N., Hironaka, K. I. and Morishita, Y. (2020). Quantitative Analysis of 3D Tissue Deformation Reveals Key Cellular Mechanism Associated with Initial Heart Looping. Cell reports 30, 3889–3903 e3885.

Kelly, R. G., Buckingham, M. E. and Moorman, A. F. (2014). Heart fields and cardiac morphogenesis. Cold Spring Harbor perspectives in medicine 4.

Kidokoro, H., Okabe, M. and Tamura, K. (2008). Time-lapse analysis reveals local asymmetrical changes in C-looping heart tube. Developmental Dynamics 237, 3545–3556.

Le Garrec, J. F., Domínguez, J. N., Desgrange, A., Ivanovitch, K. D., Raphaël, E., Bangham, J. A., Torres, M., Coen, E., Mohun, T. J. and Meilhac, S. M. (2017). A predictive model of asymmetric morphogenesis from 3D reconstructions of mouse heart looping dynamics. eLife 6.

Lopez, A. L., 3rd, Wang, S. and Larina, I. V. (2020). Embryonic Mouse Cardiodynamic OCT Imaging. J Cardiovasc Dev Dis 7.

Lorensen, W. E. and Cline, H. E. (1987). Marching cubes: A high resolution 3D surface construction algorithm. SIGGRAPH Comput. Graph. 21, 163–169.

Luo, J., Ying, K. and Bai, J. (2005). Savitzky–Golay smoothing and differentiation filter for even number data. Signal Processing 85, 1429–1434.

Madisen, L., Garner Aleena R., Shimaoka, D., Chuong, Amy S., Klapoetke Nathan C., Li, L., van der Bourg, A., Niino, Y., Egolf, L., Monetti, C., et al. (2015). Transgenic Mice for Intersectional Targeting of Neural Sensors and Effectors with High Specificity and Performance. Neuron 85, 942–958.

Meilhac, S. M. and Buckingham, M. E. (2018). The deployment of cell lineages that form the mammalian heart. Nature reviews. Cardiology 15, 705–724.

Mohun, T. J. and Anderson, R. H. (2020). 3D Anatomy of the Developing Heart: Understanding Ventricular Septation. Cold Spring Harbor perspectives in biology 12.

Muzumdar, M. D., Tasic, B., Miyamichi, K., Li, L. and Luo, L. (2007). A global double-fluorescent Cre reporter mouse. Genesis 45, 593–605.

Ocana, O. H., Coskun, H., Minguillon, C., Murawala, P., Tanaka, E. M., Galceran, J., Munoz-Chapuli, R. and Nieto, M. A. (2017). A right-handed signalling pathway drives heart looping in vertebrates. Nature 549, 86–90.

Rousseeuw, P. J. (1987). Silhouettes: A graphical aid to the interpretation and validation of cluster analysis. Journal of Computational and Applied Mathematics 20, 53–65.

Saga, Y., Miyagawa-Tomita, S., Takagi, A., Kitajima, S., Miyazaki, J. and Inoue, T. (1999). MesP1 is expressed in the heart precursor cells and required for the formation of a single heart tube. Development (Cambridge, England) 126, 3437–3447.

Schindelin, J., Arganda-Carreras, I., Frise, E., Kaynig, V., Longair, M., Pietzsch, T., Preibisch, S., Rueden, C., Saalfeld, S., Schmid, B., et al. (2012). Fiji: an open-source platform for biological-image analysis. Nature Methods 9, 676–682.

Schmidt, P., Born, J., Campen, M. and Kobbelt, L. (2019). Distortion-minimizing injective maps between surfaces. ACM Trans. Graph. 38, Article 156.

Schmidt, P., Campen, M., Born, J. and Kobbelt, L. (2020). Inter-surface maps via constant-curvature metrics. ACM Trans. Graph. 39, Article 119.

Sharpe, J. (2017). Computer modeling in developmental biology: growing today, essential tomorrow. Development (Cambridge, England) 144, 4214–4225.

Soufan, A. T., van den Berg, G., Ruijter, J. M., de Boer, P. A., van den Hoff, M. J. and Moorman, A. F. (2006). Regionalized sequence of myocardial cell growth and proliferation characterizes early chamber formation. Circ Res 99, 545–552.

Susaki, Etsuo A., Tainaka, K., Perrin, D., Kishino, F., Tawara, T., Watanabe Tomonobu M., Yokoyama, C., Onoe, H., Eguchi, M., Yamaguchi, S., et al. (2014). Whole-Brain Imaging with Single-Cell Resolution Using Chemical Cocktails and Computational Analysis. Cell 157, 726–739.

Taubin, G. (1995). Curve and surface smoothing without shrinkage. In Proceedings of IEEE International Conference on Computer Vision, pp. 852–857.

Tyser, R. C. V., Ibarra-Soria, X., McDole, K., Arcot Jayaram, S., Godwin, J., van den Brand, T. A. H., Miranda, A. M. A., Scialdone, A., Keller, P. J., Marioni, J. C., et al. (2021). Characterization of a common progenitor pool of the epicardium and myocardium. Science 371.

Voronov, D. A., Alford, P. W., Xu, G. and Taber, L. A. (2004). The role of mechanical forces in dextral rotation during cardiac looping in the chick embryo. Developmental biology 272, 339–350.

Yang, Y., Zhang, W.-X., Liu, Y., Liu, L. and Fu, X.-M. (2020). Error-bounded compatible remeshing. ACM Trans. Graph. 39, Article 113.

Yoshizawa, S., Belyaev, A. G. and Seidel, H.-P. (2003). Free-form skeleton-driven mesh deformations. In Proceedings of the eighth ACM symposium on Solid modeling and applications, pp. 247–253. Seattle, Washington, USA: Association for Computing Machinery.

Yue, Y., Zong, W., Li, X., Li, J., Zhang, Y., Wu, R., Liu, Y., Cui, J., Wang, Q., Bian, Y., et al. (2020). Long-term, in toto live imaging of cardiomyocyte behaviour during mouse ventricle chamber formation at single-cell resolution. Nature cell biology 22, 332–340.

Yushkevich, P. A., Piven, J., Hazlett, H. C., Smith, R. G., Ho, S., Gee, J. C. and Gerig, G. (2006). User-guided 3D active contour segmentation of anatomical structures: Significantly improved efficiency and reliability. NeuroImage 31, 1116–1128.

